# Functional selection in SH3-mediated activation of the PI3 kinase

**DOI:** 10.1101/2024.04.30.591319

**Authors:** Safia S. Aljedani, Abdullah Aldehaiman, Anandsukeerthi Sandholu, Siba Alharbi, Victor C.Y. Mak, Haiyan Wu, Adrien Lugari, Mariusz Jaremko, Xavier Morelli, Jonathan W. Backer, John E. Ladbury, Michał Nowakowski, Lydia W.T. Cheung, Stefan T. Arold

## Abstract

The phosphoinositide-3 kinase (PI3K), a heterodimeric enzyme, plays a pivotal role in cellular metabolism and survival. Its deregulation is associated with major human diseases, particularly cancer. The p85 regulatory subunit of PI3K binds to the catalytic p110 subunit via its C-terminal domains, stabilising it in an inhibited state. Certain Src homology 3 (SH3) domains can activate p110 by binding to the proline-rich (PR) 1 motif located at the N-terminus of p85. However, the mechanism by which this N-terminal interaction activates the C-terminally bound p110 remains elusive. Moreover, the intrinsically poor ligand selectivity of SH3 domains raises the question of how they can control PI3K. Combining structural, biophysical, and functional methods, we demonstrate that the answers to both these unknown issues are linked: PI3K-activating SH3 domains engage in additional “tertiary” interactions with the C-terminal domains of p85, thereby relieving their inhibition of p110. SH3 domains lacking these tertiary interactions may still bind to p85 but cannot activate PI3K. Thus, p85 uses a functional selection mechanism that precludes nonspecific activation rather than nonspecific binding. This separation of binding and activation may provide a general mechanism for how biological activities can be controlled by promiscuous protein-protein interaction domains.

## INTRODUCTION

Evolution is largely based on the duplication and diversification of genes (Zhang, 2003). However, gene diversification is constrained by the necessity to maintain the structure and function of the proteins they encode. As a result, proteomes can contain hundreds of protein domains with similar structure and function. This raises the question of how the interactions between these domains and their often-similar ligands can be selective (Ladbury and Arold, 2000)? This question is particularly important when a lack of selectivity can lead to severe pathologies. Here we examine a case where a ubiquitous and inherently non-selective domain, the SH3 domain, regulates a critical kinase pathway.

SH3 domains are ∼60-80 amino acid protein-protein interaction domains that bind to proline-rich (PR) sequences, often containing the core PXXP motif (Ladbury and Arold, 2011; Li, 2005)(**Figure 1A**). The human proteome encodes more than 300 different SH3 domains. PR motifs are abundant in the human proteome and do not need post-translational modifications for binding. Most SH3 domains display moderate to low affinities towards PR motifs (with dissociation constants, *Kd*s, of 1–100 µM) and cross-react with different PR motifs (Ladbury and Arold, 2011). As a result, the difference in affinity of SH3 domains between their biologically relevant PR motifs, and non-specific PR motifs appears too small to exclude cross-reactivity and erroneous binding events (Ladbury and Arold, 2000, 2012; Li, 2005). Nonetheless, many SH3 domain-containing proteins contribute to highly specific signalling pathways, many of which involve kinases.

**Figure 1.**
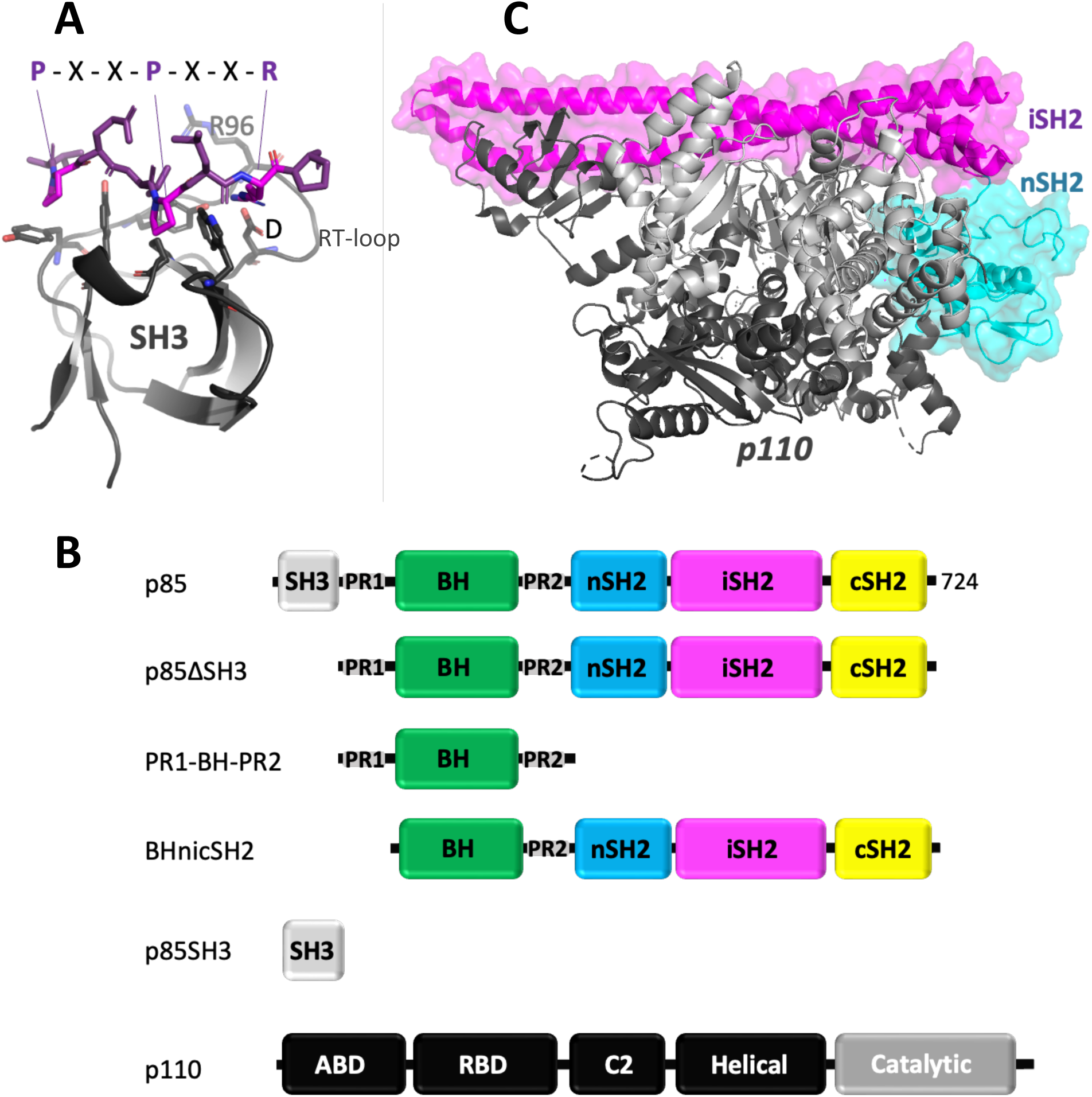
Structure and interactions of p85, p110 and SH3. **A**) Example of a canonical interaction between the Fyn SH3 domain (black ribbon) and a PXXP containing peptide (stick model with carbon atoms in magenta). The PR motif inserts its two key prolines (pink) into hydrophobic grooves on the SH3 domain (residues shown stick models). In the PR motif, the position of an arginine with respect to the prolines determines the binding orientation by engaging a salt bridge with an aspartic acid from the SH3 RT loop (Feng et al., 1994) Shown is a class II peptide (RXXPXXP) bound to Fyn SH3 (extracted from the HIV-1 Nef:Fyn SH3 complex, PDB 1AVZ). R96 of Fyn SH3 is shown as stick model. **B**) Top: Schematic drawing of full-length p85 and relevant fragments thereof used in this work. Bottom: schematic drawing of p110. The colours are maintained throughout the manuscript. **C**) Structure of p110 (black, with catalytic domain in grey) kept in an inhibited conformation through interactions with p85 iSH2 (pink) and nSH2 (cyan). Structure taken from PDB 8DCP.

Humans encode more than 500 kinase domains (Milanesi et al., 2005). The phospho-transfer activity of kinases generally triggers signalling events that can lead to transformative responses in the cell. Consequently, kinase activation is tightly controlled, and kinase deregulation is associated with severe pathologies. The phosphoinositide 3 kinases (PI3Ks) phosphorylate the 3 position hydroxyl group of phosphatidylinositol in response to growth factors and environmental cues. PI3K activation triggers cell growth, proliferation, differentiation, motility survival and intracellular trafficking (Backer, 2010; Engelman et al., 2006). Erroneous activation of the PI3K pathway is a leading cause of cancer and is associated with neurodegenerative disorders (Rai et al., 2019; Vanhaesebroeck et al., 2010).

The class IA PI3K, comprising α, β and γ isoforms, are heterodimers formed by a p85 regulatory subunit and a p110 catalytic subunit. Upon binding, p85 stabilises p110 against degradation, but keeps it in a catalytically inactive form (Backer, 2010; Yu et al., 1998b). p85 is composed of a Src homology (SH) 3 domain, a RhoGap/breakpoint cluster region homology (BH) domain, and two SH2 domains (nSH2 and cSH2) separated by a coiled-coil inter-SH2 domain (iSH2)(**Figure 1B**). The p85 iSH2 domain associates with p110, stabilising it. In the absence of activating ligands, additional interactions with the p85 SH2 domains inhibit the catalytic activity of p110 (Hale et al., 2009; Mandelker et al., 2009; Yu et al., 1998b, 1998a; Zhang, 2003) (**Figure 1B**). These interactions and their dynamics show some variability among the class IA PI3Ks. For PI3Kα, the minimal regulatory fragment is composed of the p85 nSH2 and iSH2 domains (Yu et al., 1998a). Structural data revealed that the nSH2 domain makes inhibitory contacts with the helical, C2 and kinase domains of p110α (Huang et al., 2007; Mandelker et al., 2009). In the case of the PI3Kβ heterocomplex, structural analysis suggests that the cSH2 domain binds directly to the kinase domain (Hart et al., 2022). Although this interaction with the cSH2 has not been observed in PI3Kα, the p85α cSH2 domain does appear to regulate kinase activation in PI3Kα (Jücker et al., 2002; Yu et al., 1998a) and has been experimentally shown in the complex to reside close to the active site of the p110α kinase domain. However, in PI3Kα the cSH2 domain appears to be more flexibly localised in this position than in PI3Kβ (Hart et al., 2022). Collectively, these structural observations explain how ligands that bind to p85α SH2 domains (i.e. ligands containing phosphotyrosine motifs) can activate p110α by disrupting the inhibitory SH2:p110α interactions. This mechanism also explains the disease-associated effects of many mutations in these domains (Carpenter et al., 1993; Li et al., 2021). However, this model cannot explain how ligands of the other p85α regions activate p110α. Here we investigate how SH3 domains can activate p110α by binding to the proline-rich region 1 (PR1) of p85α.

It has been known for more than 30 years that the SH3 domains from several Src family protein tyrosine kinases (i.e., Lyn, Lck, Fyn, Fgr, and Hck) can activate PI3K by binding to PR1, situated between the p85α SH3 domain and the BH domain (Axelsson et al., 2000; Kapeller et al., 1994; Pleiman et al., 1994; Prasad et al., 1993; Renzoni et al., 1996) (**Figure 1B**). Intriguingly, the PR1 region is more than 250 residues away in the amino acid sequence from the nSH2 domain.

Furthermore, cryo-EM structures of p85α bound to p110α lack clear density for the SH3-PR1-BH domains (Hart et al., 2022; Liu et al., 2021), inferring that these domains are not stably associated within the complex formed between p110α and p85α. Hence, it is unknown how SH3 domains can mechanistically activate p110α, and how such an activation based on promiscuous SH3 domain interactions can be selective. Resolving these longstanding questions will clarify the regulation of PI3K pathways in normal and diseased states, and, more generally, help understanding how cellular specificity can be achieved with intrinsically non-specific binding modules. Here, we combine biophysical and functional assays to show that SH3 domains achieve specific PI3K activation through a functional selection mechanism.

## RESULTS

### Tertiary interactions affect the association between SH3 domains and p85

We chose the Fyn SH3 domain to investigate its activation of PI3K. Renzoni and colleagues previously studied the binding of Fyn SH3 to p85α (for simplicity, we hereafter refer to p85α and p110α as p85 and p110, respectively). They observed that the almost complete p85 containing PR1 (p85ΔSH3; the N-terminal p85 SH3 domain was deleted to avoid effects from intramolecular competition) bound to the Fyn SH3 domain with a higher affinity and different thermodynamic parameters than did a peptide derivative of PR1 (Renzoni et al., 1996). It was hypothesised that the differences resulted from p85 interactions outside the canonical class I PXXP motif (<u>R</u>^93^PL<u>P</u>VA<u>P</u>). Indeed, our ITC experiments with Fyn SH3 and either a PR1 peptide or p85ΔSH3 confirmed that Fyn SH3 bound p85ΔSH3 with a significantly higher affinity and a markedly different thermodynamic profile compared to the PR1 peptide (see entries in light grey in **Table 1; Supplementary Figure 1**).

**Table 1:**
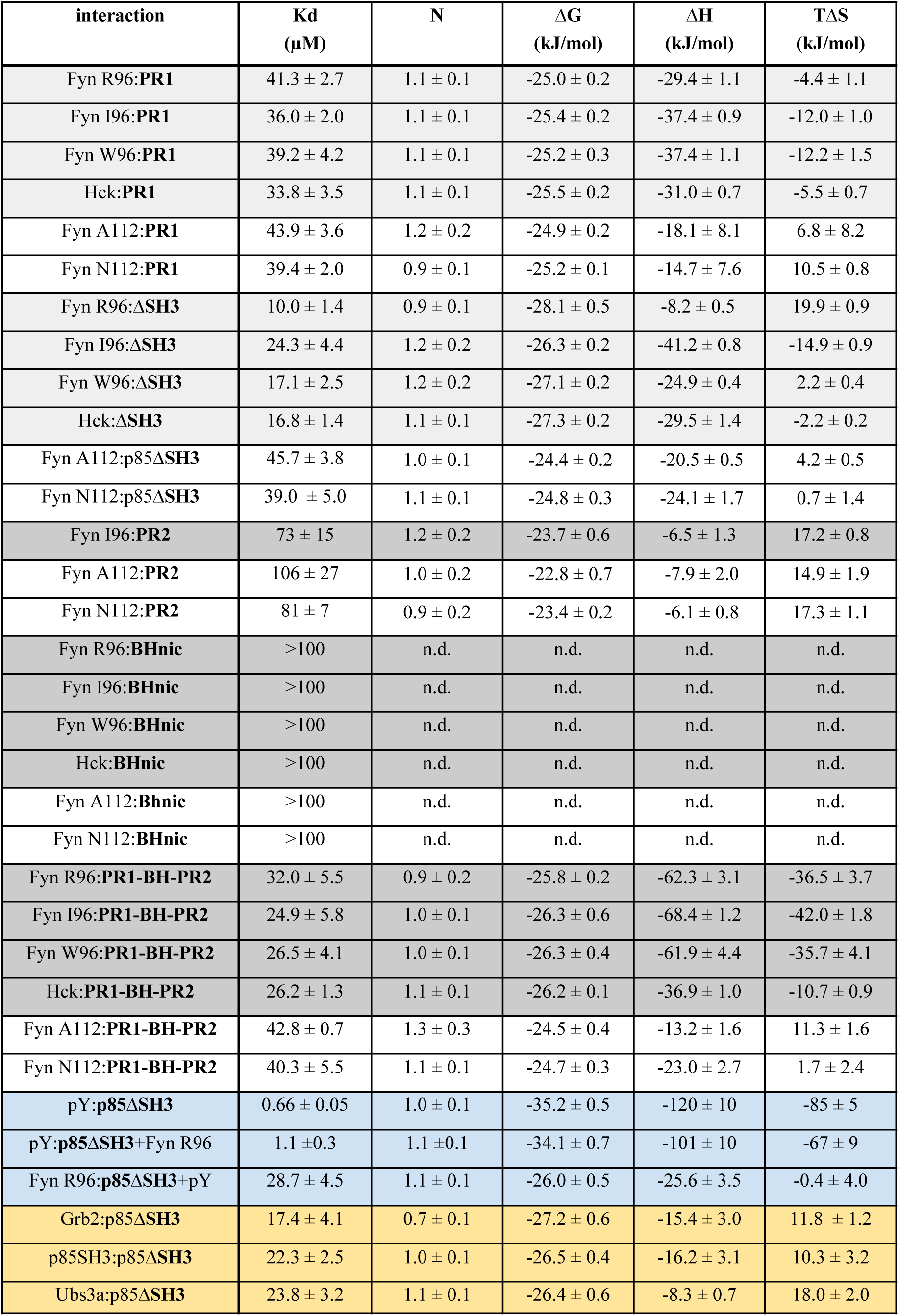
Summary of thermodynamic parameters. . Fields are coloured according to types of interaction, to facilitate associating data with the manuscript text. ITC data are shown in Supplementary Figure 1. Errors are based on triplicate repeats.

| **interaction** | **Kd**<br>**(μM)** | **N** | **ΔG**<br>**(kJ/mol)** | **ΔH**<br>**(kJ/mol)** | **TΔS**<br>**(kJ/mol)** |
| --- | --- | --- | --- | --- | --- |
| Fyn R96:PR1 | 41.3 ± 2.7 | 1.1 ± 0.1 | -25.0 ± 0.2 | -29.4 ± 1.1 | -4.4 ± 1.1 |
| Fyn I96:PR1 | 36.0 ± 2.0 | 1.1 ± 0.1 | -25.4 ± 0.2 | -37.4 ± 0.9 | -12.0 ± 1.0 |
| Fyn W96:PR1 | 39.2 ± 4.2 | 1.1 ± 0.1 | -25.2 ± 0.3 | -37.4 ± 1.1 | -12.2 ± 1.5 |
| Hck:PR1 | 33.8 ± 3.5 | 1.1 ± 0.1 | -25.5 ± 0.2 | -31.0 ± 0.7 | -5.5 ± 0.7 |
| Fyn A112:PR1 | 43.9 ± 3.6 | 1.2 ± 0.2 | -24.9 ± 0.2 | -18.1 ± 8.1 | 6.8 ± 8.2 |
| Fyn N112:PR1 | 39.4 ± 2.0 | 0.9 ± 0.1 | -25.2 ± 0.1 | -14.7 ± 7.6 | 10.5 ± 0.8 |
| Fyn R96:ΔSH3 | 10.0 ± 1.4 | 0.9 ± 0.1 | -28.1 ± 0.5 | -8.2 ± 0.5 | 19.9 ± 0.9 |
| Fyn I96:ΔSH3 | 24.3 ± 4.4 | 1.2 ± 0.2 | -26.3 ± 0.2 | -41.2 ± 0.8 | -14.9 ± 0.9 |
| Fyn W96:ΔSH3 | 17.1 ± 2.5 | 1.2 ± 0.2 | -27.1 ± 0.2 | -24.9 ± 0.4 | 2.2 ± 0.4 |
| Hck:ΔSH3 | 16.8 ± 1.4 | 1.1 ± 0.1 | -27.3 ± 0.2 | -29.5 ± 1.4 | -2.2 ± 0.2 |
| Fyn A112:p85ΔSH3 | 45.7 ± 3.8 | 1.0 ± 0.1 | -24.4 ± 0.2 | -20.5 ± 0.5 | 4.2 ± 0.5 |
| Fyn N112:p85ΔSH3 | 39.0 ± 5.0 | 1.1 ± 0.1 | -24.8 ± 0.3 | -24.1 ± 1.7 | 0.7 ± 1.4 |
| Fyn I96:PR2 | 73 ± 15 | 1.2 ± 0.2 | -23.7 ± 0.6 | -6.5 ± 1.3 | 17.2 ± 0.8 |
| Fyn A112:PR2 | 106 ± 27 | 1.0 ± 0.2 | -22.8 ± 0.7 | -7.9 ± 2.0 | 14.9 ± 1.9 |
| Fyn N112:PR2 | 81 ± 7 | 0.9 ± 0.2 | -23.4 ± 0.2 | -6.1 ± 0.8 | 17.3 ± 1.1 |
| Fyn R96:BHnic | >100 | n.d. | n.d. | n.d. | n.d. |
| Fyn I96:BHnic | >100 | n.d. | n.d. | n.d. | n.d. |
| Fyn W96:BHnic | >100 | n.d. | n.d. | n.d. | n.d. |
| Hck:BHnic | >100 | n.d. | n.d. | n.d. | n.d. |
| Fyn A112:Bhnic | >100 | n.d. | n.d. | n.d. | n.d. |
| Fyn N112:BHnic | >100 | n.d. | n.d. | n.d. | n.d. |
| Fyn R96:PR1-BH-PR2 | 32.0 ± 5.5 | 0.9 ± 0.2 | -25.8 ± 0.2 | -62.3 ± 3.1 | -36.5 ± 3.7 |
| Fyn I96:PR1-BH-PR2 | 24.9 ± 5.8 | 1.0 ± 0.1 | -26.3 ± 0.6 | -68.4 ± 1.2 | -42.0 ± 1.8 |
| Fyn W96:PR1-BH-PR2 | 26.5 ± 4.1 | 1.0 ± 0.1 | -26.3 ± 0.4 | -61.9 ± 4.4 | -35.7 ± 4.1 |
| Hck:PR1-BH-PR2 | 26.2 ± 1.3 | 1.1 ± 0.1 | -26.2 ± 0.1 | -36.9 ± 1.0 | -10.7 ± 0.9 |
| Fyn A112:PR1-BH-PR2 | 42.8 ± 0.7 | 1.3 ± 0.3 | -24.5 ± 0.4 | -13.2 ± 1.6 | 11.3 ± 1.6 |
| Fyn N112:PR1-BH-PR2 | 40.3 ± 5.5 | 1.1 ± 0.1 | -24.7 ± 0.3 | -23.0 ± 2.7 | 1.7 ± 2.4 |
| pY:p85ΔSH3 | 0.66 ± 0.05 | 1.0 ± 0.1 | -35.2 ± 0.5 | -120 ± 10 | -85 ± 5 |
| pY:p85ΔSH3+Fyn R96 | 1.1 ± 0.3 | 1.1 ± 0.1 | -34.1 ± 0.7 | -101 ± 10 | -67 ± 9 |
| Fyn R96:p85ΔSH3+pY | 28.7 ± 4.5 | 1.1 ± 0.1 | -26.0 ± 0.5 | -25.6 ± 3.5 | -0.4 ± 4.0 |
| Grb2:p85ΔSH3 | 17.4 ± 4.1 | 0.7 ± 0.1 | -27.2 ± 0.6 | -15.4 ± 3.0 | 11.8 ± 1.2 |
| p85SH3:p85ΔSH3 | 22.3 ± 2.5 | 1.0 ± 0.1 | -26.5 ± 0.4 | -16.2 ± 3.1 | 10.3 ± 3.2 |
| Ubs3a:p85ΔSH3 | 23.8 ± 3.2 | 1.1 ± 0.1 | -26.4 ± 0.6 | -8.3 ± 0.7 | 18.0 ± 2.0 |

To identify the molecular basis for the difference in binding, we took inspiration from previous analyses that demonstrated that the Nef proteins from simian and human immunodeficiency viruses (SIV and HIV, respectively) select SH3 domains by recognizing surfaces beyond the canonical PR binding site. A key feature of this tertiary recognition is a pocket on Nef that binds to the residue in position “R” of the so-called SH3 RT loop (residue R96 in Fyn SH3 domain). In HIV-1 Nef, this pocket is hydrophobic and, therefore, favours an interaction with SH3 domains that have a hydrophobic residue in the “R” position, such as Hck with an isoleucine, or the Fyn R96I and R96W mutants (referred to as I96 and W96 in this manuscript)(Arold et al., 1997; Collette et al., 2000; Horenkamp et al., 2011; Lee et al., 1996, 1995). However, the tertiary association also rearranges and fixes the SH3 RT loop onto the Nef surface, favouring SH3 domains with intrinsically more flexible RT loops (case of Hck and Fyn W96) (Aldehaiman et al., 2021; Arold et al., 1998). Thus, the stereochemical match of this SH3 residue and the Nef pocket as well as the effect of the position 96 on the RT loop flexibility control the affinity and selectivity of Nef toward SH3 domains. To test if the SH3 “R” position also affects SH3:p85 interactions, we used ITC to study the interaction between p85 and the Fyn R96, I96, and W96 SH3 domains. Fyn R96, I96 and W96 bound to PR1 with comparable medium-to-low affinities (with dissociation constants, *Kd*s, of approximately 40 µM under our conditions) and similar enthalpy-dominated energy changes [**Table 1** (light grey)**, Supplementary Figure 1**]. Conversely, these SH3 domains showed a more than twofold difference in their *Kd*s to p85ΔSH3, resulting from markedly changed enthalpy and entropy contributions. However, in contrast to the Nef:SH3 interactions, Fyn R96 displayed the highest affinity (*Kd* of 10.0 ± 1.4 µM), whilst Fyn W96 bound with an almost twofold weaker *Kd* to p85ΔSH3. We concluded that the SH3 RT loop residues modulate the association with p85ΔSH3, but not with PR1, and that the presence of additional domains of the p85ΔSH3 construct affects the association of SH3 with p85 PR1. However, the effect of additional SH3 regions differs between p85ΔSH3 and Nef.

### The C-terminal p85 domains affect the association with SH3 domains

We next used ITC to help identify the domains of p85 responsible for the tertiary effect. The impact of the PR2 motif could be disregarded because it bound with negligible affinity (*Kd* >70 µM) to the SH3 domains, and this residual binding was further diminished in the BH-PR2-nicSH2 construct [**Table 1** (dark grey)**, Supplementary Figure 1**]. The observation that the PR2 motif does not play a significant role in the binding to SH3 domains was in agreement with previous observations (Pleiman et al., 1994). Binding experiments with the p85 PR1-BH-PR2 fragment did not recapitulate the affinity and SH3 variant–specific thermodynamics observed for p85ΔSH3 [**Table 1** (dark grey)**, Supplementary Figure 1**], implying that the C-terminal nSH2, iSH2, and cSH2 domains contributed to the binding characteristics of p85ΔSH3. This potential involvement of the additional domains at the C-terminal of p85 domains was interesting, as these domains are responsible for binding and inhibiting p110 (Backer, 2010; Yu et al., 1998a).

To explore the possibility that Fyn SH3 binding engages the C-terminal part of p85, we tested the influence of a platelet-derived growth factor receptor (PDGFR) peptide known to bind to both p85 nSH2 and cSH2 domains (Klippel et al., 1992). This peptide (referred to as pY) comprised PDGFR residues 738-755, with phosphorylated tyrosines in position 740 and 750. We observed that adding pY into the ITC cell containing p85ΔSH3 lowered the affinity of Fyn R96 for p85ΔSH3 and changed the enthalpy and entropy contributions. The presence of Fyn R96 in the measurement cell only marginally lowered the affinity for pY when titrated onto p85ΔSH3, in line with the affinity of p85ΔSH3 bing 15-fold higher for pY than for Fyn R93 [**Table 1** (light blue), **Supplementary Figure 1**]. This result supports that pY and Fyn R96 bind p85ΔSH3 on at least partially overlapping regions. Considering that pY binds to the SH2 domains of p85, but not to the PR1 region, one or both SH2 domains appear to be involved in the interaction with Fyn SH3.

### PI3K activation does not correlate with SH3 affinity

From the above experiments, we have shown that the interaction of p85 with the Fyn SH3 domain variants requires an interface that extends to include the C-terminal region. Since this region is known to regulate the catalytic activity of the p110 subunit, we performed lipid kinase assays to assess the impact of SH3 binding. These assays were performed based on p110 immunoprecipitated from endometrial cancer cells (EFE-184) which carry wild-type p110 and p85 (see **Methods**).

As a positive control, we used the PDGFR pY phosphopeptide. Two and 10 µM of pY peptide activated PI3K in a dose-dependent manner (**Figure 2A**), in line with the *Kd* of approximately 660 nM for the association between pY and p85ΔSH3 [**Table 1** (light blue), **Supplementary Figure 1**]. 100 µM Fyn SH3 R96, I96 and W96 also activated PI3K, although with lower efficacy than the pY peptide at 10 µM. The activation efficacy differed among the three SH3 domains, with Fyn W96 > Fyn I96 > Fyn R96. Remarkably, this difference in efficacy, caused by a mutation that is not directly involved in PR1 binding, did not correlate with the *in vitro* binding affinities (i.e., Fyn R96 > W96 > I96).

**Figure 2.**
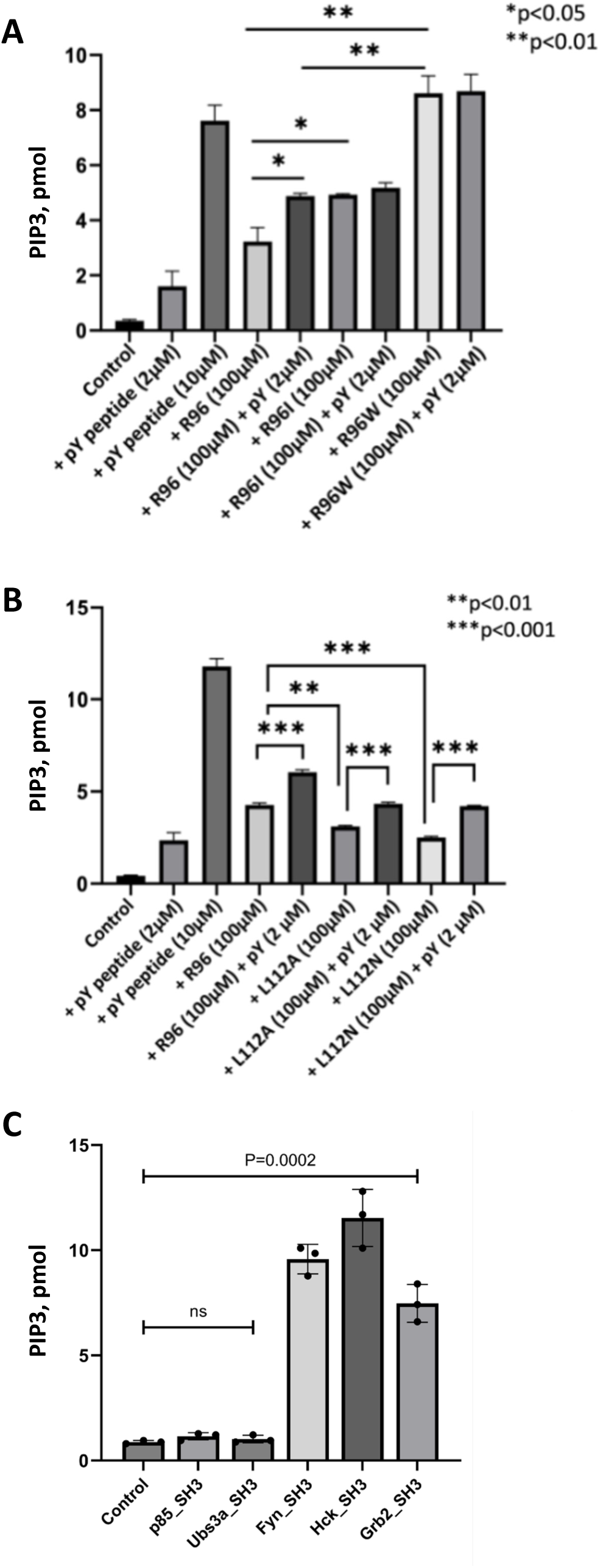
Lipid kinase assays. Immunoprecipitated p110α protein from EFE-184 cells was incubated with **A**) doubly phosphorylated PDGFR peptide (pY) and/or recombinant purified wild-type Fyn (R96) or SH3 mutant (R96I or R96W), **B**) pY and/or recombinant purified wild-type Fyn (R96) or SH3 mutant L112A or L112N (with wild-type R96 background), or **C**) wild-type SH3 domain of the indicated proteins (100 µm), prior to lipid kinase assay and absorbance measurement. Reaction without incubation with any peptide served as control. Immunoprecipitation using normal IgG instead of anti-p110α antibody in the control sample was performed in parallel for background signal. Data in graphs are levels of PIP3 after subtracting the absorbance of IgG samples and are presented as the mean of triplicates ± S.D. P-values were obtained using unpaired t-test. ns, not statistically significant.

Adding 2 µM pY peptide at the same time as the SH3 domains increased PI3K activation by Fyn R96, but not by Fyn I96 or W96 (**Figure 2A**). The PI3K activation by Fyn I96 or W96 alone was already on par or higher compared to the combined activation by Fyn R96 and the 2 µM pY peptide. We hypothesised that, compared to Fyn R96, the Fyn I96 or W96 variants more effectively engaged with p85ΔSH3 regions that overlap with pY activating interactions, eliminating the effect of adding pY.

### Interface mapping identifies additional site for complex formation between Fyn SH3 and p85

To investigate the molecular basis for the observed effects, we first titrated the Fyn SH3 domains Fyn R96, I96 and W96 with increasing amounts of PR1. For all three SH3 domain variants, SH3:PR1 ratios of 1:0.5, 1:1, 1:2, 1:4, 1:8, 1:16 were measured under the same conditions. In these titrations, we observed significant chemical shift changes at the SH3:PR1 ratio of 1:0.5 (W96 and I96) or 1:1 (R96) (**Figure 3A, Supplementary Figure 2**). Methyl groups of isoleucines changed their chemical shifts in titrations of all three constructs, with an intermediate regime for Fyn W96 and I96, and a fast exchange for R96. The chemical shift changes for Fyn R96, I96 and W96 upon binding to PR1 were very similar, and in agreement with the imprint expected from a canonical PR motif (**Figure 3A, Supplementary Figure 2**). We concluded that the peptide bound in the same canonical way to all three SH3 variants tested.

**Figure 3.**
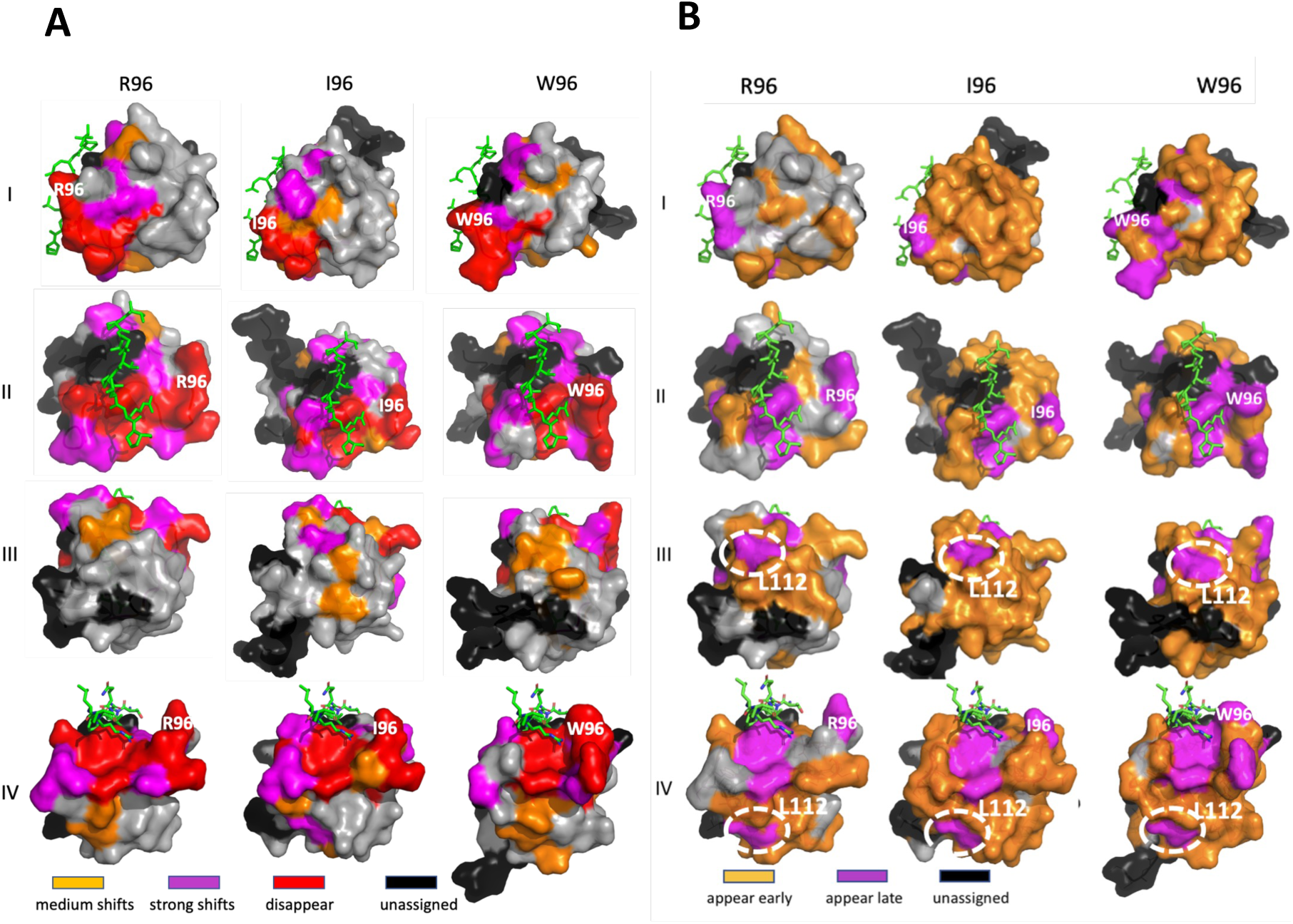
NMR interaction mapping onto Fyn SH3 variants. The crystal structure of Fyn R96 (PDB accession 1AVZ), Fyn I96 (6IPY), and Fyn W96 (6IPZ) were used to map the chemical shifts. The PR sequence from the HIV-1 Nef protein (PDB 1AVZ; shown as green stick model) has been taken as a guide for ‘canonical’ binding of a PR to Fyn SH3. Four different orientations are shown (I-IV). **A**) Binding of the PR1 motif to Fyn SH3 wild type (R96) and the I96 and W96 mutants, as mapped by chemical shift analysis. **B**) Binding of 15N labelled Fyn SH3 R96, I96 and W96 to p85ΔSH3. Shown are the residues that reappear early during the titration (at p85ΔSH3: SH3 ratios of 1:4, 1:3 and 1:6 for WT, RI and WR, respectively) or later during the titration (at p85ΔSH3:SH3 ratios of 1:6, 1:6 and 1:8 for WT, RI and WR, respectively).

We next investigated the binding of the Fyn SH3 variants to p85ΔSH3. Due to the limited solubility of p85ΔSH3, we performed ^1^H-^15^N HSQC titrations where we titrated increasing amounts of ^15^N-labelled SH3 domains onto unlabelled p85ΔSH3 kept at a constant concentration of 100 µM. Ratios of p85ΔSH3:SH3 domains were 1:0.1, 1:0.25, 1:0.5, 1:1, 1:2, 1:3, 1:4, 1:6, 1:8, 1:10, 1:15, and 1:25. For the analysis it is important that the PR1 motif is separated from the BH domain by 15 residues which are predicted to be flexible from their primary sequence, AlphaFold model (pLDDT <70 in <u>alphafold.ebi.ac.uk/entry/P27986</u>), and small angle X-ray scattering analysis (**Supplementary Figure 3**). If there are no additional interactions between the SH3 domains and p85ΔSH3 beyond the linear PR1 region, then the NMR signals for p85ΔSH3-bound SH3 domains would resemble those we recorded for the titrations with the PR1 peptide alone. Conversely, if there are additional tertiary interactions that fix the bound SH3 domains onto one or more domains of p85ΔSH3, then chemical shifts of the labelled SH3 domain would broaden relative to the size and stability of the complex formed.

Our experimental observations clearly demonstrate the second case. At the highest p85ΔSH3:SH3 ratios (1:0.1), most chemical shifts disappeared in all three mutants, inferring the firm engagement of the SH3 domain in a large complex (**Figure 3B**). Only visible (and hence highly dynamic; not restricted by the complex) were residues V84, D100 and L101 for Fyn R96 and D78 and T85 for Fyn W96. As the p85ΔSH3:SH3 ratios diminished, gradually more and more residues reappeared. For low SH3 concentrations, we observed peaks for particular (flexible) residues that were shifted from their position in free SH3, which suggested a fast exchange regime. At p85ΔSH3:SH3 ratios of 1:6 (Fyn R96 and I96) or 1:8 (Fyn W96) most peaks reappear. However, some residues remained invisible and only reappeared later in the titrations, indicating these as the strongest bound residues (**Figure 3B**). Position 96 of Fyn wild-type and mutants was among the strongly bound residues, confirming the importance of this position for the association with p85ΔSH3. This NMR experiment also identified Fyn L112 as strongly bound, even though it is remote from the PR binding surface.

Together, these results demonstrated that Fyn SH3 is not flexibly attached by canonical linear interactions in the complex with p85ΔSH3 but is fixed to one or more domains through supplementary interactions involving Fyn positions R96, located in the RT loop close to the canonical PR binding site (‘PR-proximal’), and L112 on the opposite site of the SH3 domain (‘PR-distal’).

### SH3 L112 mutants impair PI3K activation

Our NMR analysis suggested that the PR-distal L112 position contributes to the formation of a ‘heavy’ multidomain complex with p85. To test the functional repercussions of mutating this position, we generated the L112N and L112A mutations based on the wild-type Fyn R96 SH3 domain. ITC titrations showed that these L112 mutant SH3 domains bound the p85ΔSH3 with the same affinity (*Kd* ∼40 µM) and comparable ΔH and TΔS contributions as the PR1 peptide [**Table 1** (white)**, Supplementary Figure 1**]. Our differential scanning fluorimetry (DSF) measurements confirmed that the Fyn SH3 L112N and L112A variants had similar stability as the wild-type domain (i.e., *Tm* above 70° C), and hence that altered p85 binding was not due to SH3 destabilisation **(Supplementary Figure 4**). These observations corroborated that the PR-distal mutations directly affected Fyn SH3 associations with p85ΔSH3.

In our lipid kinase assay, the L112N and L112A mutations significantly decreased the capability of the Fyn SH3 domain to activate PI3K (**Figure 2B**). As observed for the wild-type Fyn SH3 domain, the addition of the pY peptide increased the PI3K activation by L112N and L112A mutant SH3 domains. This additional activation by pY was roughly constant, hence the combined activation level promoted by pY and mutants together remained lower than the combined activation of pY and wild-type Fyn SH3. We concluded that the NMR-identified L112 position of Fyn was required to achieve the tertiary effect on p85 binding, and the functional effect on PI3K activation.

### Predicting PI3K activation of SH3 domains based on the tertiary site

Having demonstrated the importance of Fyn L112 for activating PI3K, we asked whether it would be possible to predict the PI3K activating potential of other SH3 domains, based on this region. A medium-sized hydrophobic residue is also found in the SH3 domain of the Src family kinase Hck (corresponding to L108), and in the C-terminal SH3 domain (cSH3) of the Grb2 adaptor protein (M186). Hence, we predicted that these domains bind and activate PI3K. Conversely, p85’s own SH3 domain and the SH3 domain of Ubs3a have this PR1-remote tertiary region completely altered in charge (polar) and in shape (through the insertion of a helix) (**Figure 4**). Therefore, we predicted that these domains cannot engage in the requited tertiary contacts with p85 to activate PI3K, even though they may bind to PR1.

**Figure 4.**
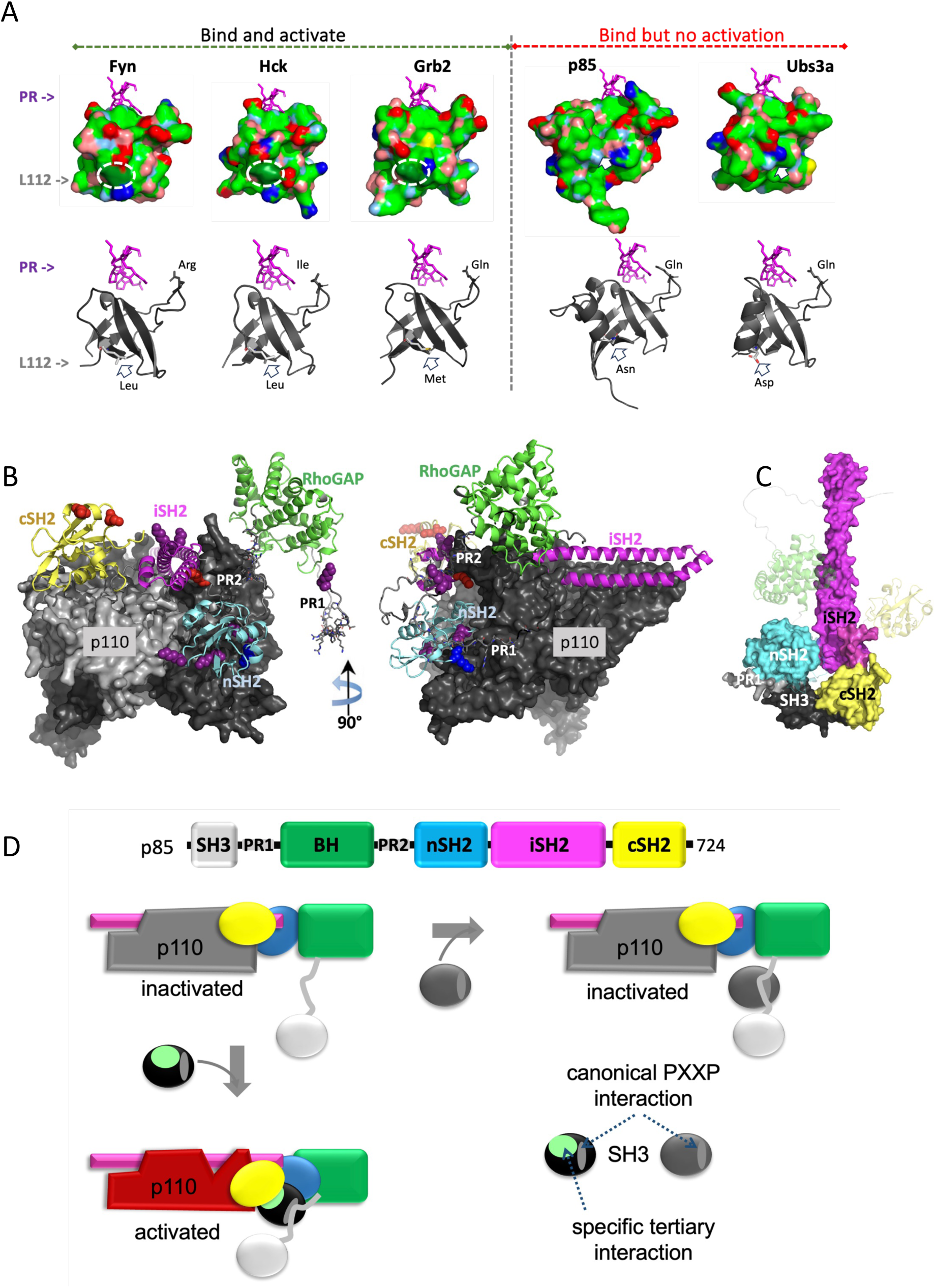
Molecular mechanism for the activation of p85 by SH3 domains. **A**) Classification of SH3 domains that either bind and activate p85 (left) or just bind to p85 without activating it (right). *Top*: SH3 domain surfaces, colour coded according to atoms being hydrophobic: green; sulphur: yellow; acidic: red; basic: blue; polar nitrogen: cyan; polar-oxygen: salmon. The PR sequence from the HIV-1 Nef protein (PDB 1AVZ; shown as magenta stick model) has been taken as a guide for ‘canonical’ binding of a PR to SH3 domains. The position corresponding to Fyn L112 is circled in ‘activating’ SH3 domains, and roughly indicated by an arrow in ‘non-activating’ SH3 domains, where the structure at this position is different. *Bottom*: Ribbon model corresponding to the surface view. The PR motif is in magenta. L112 position and nature of corresponding residue is given. **B**) Two 90° views XLMS-mapped lysines. p110 is shown as a surface model with the catalytic domain in light grey, and other domains in dark grey. Domains of p85ΔSH3 are colour-coded. Lysines identified in XLMS and being compatible with a membrane-associated p110:p85 complex are shown as sphere models, coloured in blue (crosslinked only in presence of Fyn SH3), purple (only Grb2 cSH3), or red (crosslinked with both Fyn and Grb2 SH3). The p110:p85ΔSH3 model was produced by AlphaFold. **C**) Speculative 3D model to illustrate how binding of an activating SH3 domain may contact and rearrange both SH2 domains of p85. Model produced by AlphaFold. **D**) Speculative model for PI3K activation through SH3 domains. The schematic p85 drawing recapitulates the domain colours. p110 is shown in dark grey. *Top left*: In absence of stimulating ligands, p85 binds p110 through the iSH2 domain. Interactions between the p85 SH2 domains and p110 keep p110 in the catalytically inactive conformation. *Top right*: SH3 domains that lack the tertiary interfaces may still bind to p85 PR, but cannot activate PI3K. *Bottom left*: When an SH3 domain with correct tertiary surface binds (green; only one such surface is shown), then the resulting interactions with the p85 nSH2-iSH2-cSH2 fragment remove the inhibitory SH2 interactions, activating p110 (red). *Bottom right*: Schematic drawing of SH3 domains with or without tertiary surfaces.

Using ITC, we showed that all SH3 domains were able to bind to p85ΔSH3 within a limited range of affinities (*Kd*s of 7–24 µM), comparable to those of Fyn I96 and W96 [**Table 1** (yellow)**, Supplementary Figure 1**]. In our lipid kinase assays, Grb2 cSH3 and Hck SH3 activated PI3K similarly to Fyn SH3 (**Figure 2B**). Despite their similar affinity to p85ΔSH3, the SH3 domains of p85 and Ubs3a failed to activate PI3K. We concluded that the stereochemistry of the PR-distal tertiary sites, but not the binding affinity, correlated with the capability of these SH3 domains to activate PI3K.

### Crosslinking Mass Spectrometry (XL-MS) suggests SH3 interactions with p85 domains

To further clarify the mechanism by which SH3 domains activate PI3K, we crosslinked the SH3 domains from Fyn, Grb2, and p85 with p85ΔSH3 using bissulfosuccinimidyl suberate (BS3). Crosslinked proteins were separated on SDS-PAGE gel, trypsinised, and analysed by MS (**Supplementary Figure 5**). In addition to the N-terminus, BS3 crosslinks amide groups from lysines, of which the SH3 domains of Fyn, Grb2, and p85 have two, three and three, respectively. XL-MS resulted in eight different crosslinks for Fyn, twelve for Grb2 and 21 for p85 SH3 (**Supplementary Figure 6** A-C). The SH3 domains of Fyn and Grb2 showed a similar crosslinking pattern involving lysines in the p85 nSH2, iSH2 and cSH2 domains. Grb2 cSH3 also crosslinked to the p85ΔSH3 N-terminal amide, seven residues upstream of PR1. p85 SH3 showed crosslinks with all domains, including the BH domain, and especially in the iSH2 domain (12 crosslinks, compared to five and six crosslinks for Fyn and Grb2, respectively). Thus, the PI3K-activating SH3 domains from Fyn and Grb2 showed a more restricted interaction pattern in XL-MS than the non-activating p85 SH3. This difference in the XL-MS pattern may result from the different positions of the lysines on these SH3 domains. However, it may also infer a larger mobility and accessibility of the p85 SH3 lysines that allow more crosslinks with the sticky hydrophobic patches that iSH2 uses to bind to p110 or tether the coiled-coil structure. Conversely, the cross-linking potential of Fyn and Grb2 SH3 domains may be comparatively restricted by more stable tertiary interactions with the SH2 domains.

### Proposed mechanistic model for SH3-mediated PI3K activation

Our efforts to produce a crystal structure of SH3 domains bound to p85 (and fragments thereof) failed, likely due to the large flexibility of this protein. Atomic structure determination also appeared unfeasible with NMR (due to the observed peak broadening) or cryo-EM (due to the low SH3 affinity and the persistent flexibility of the PR1, BH, PR2 and nSH2 domains even in a cross-linked, nanobody-stabilised complex with p110; (Hart et al., 2022). Therefore, to produce a mechanistic model for the SH3-promoted activation of PI3K, we projected the crosslinks onto the biological p85:p110 heterocomplex (**Figure 4B**). Because an experimental all-domain structure of PI3K is not available, we generated a model of this complex with AlphaFold (’Guzmán-Vega and ’Arold, 2024; Jumper et al., 2021). The predicted p85:p110 complex superimposed closely with experimental models for p85:p110, also reproducing the experimentally observed positions of p85 nSH2 and iSH2 (**Supplementary Figure 7A**,B). In this AlphaFold model, the cSH2 domain is positioned next to the kinase domain. This position is similar to the inhibitory cSH2 position identified in the crystal structure of the PI3β heterocomplex (PDB id 2Y3A) (Zhang et al., 2011). However, unlike the PI3Kβ crystal structure, the cSH2 of the PI3Kα AlphaFold model was not in direct contact with the p110 kinase domain, in line with the lack of density for this domain in experimental PI3Kα structures (Hart et al., 2022). Rather, cSH2 was flexibly anchored through a predicted interaction between the iSH2-cSH2 linker residues Y^607^SLV and the p110α kinase domain, suggestive of a different inhibitory mechanism involving Y^607^SLV (**Supplementary Figure 7C**). The p85 BH and PR1 regions were not predicted to stably associate with the rest of the complex, in line with previous reports (Hart et al., 2022). We used this p85:p110 model to eliminate those lysines from the XL-MS data that would not be available for SH3 interactions in the biological context, because of being occluded by either a stable p110 association or the membrane. This constraint eliminated most lysines on iSH2, except for those at the basis of its coiled-coil structure (**Figure 4B**). Consequently, our data suggested that the activating SH3 domains interact with the nSH2, cSH2 and the basis of the iSH2 in addition to the PR1. Considering that the Fyn SH3 positions 96 and 112 that were important for binding and activation are located on different sides of the SH3 domain, it is conceivable that activation results from these positions contacting residues on the nSH2, iSH2 and cSH2 domains of p85. Hence, a possibility in line with our data would be that upon binding to PR1, activating SH3 domains form an adaptor that simultaneously engages the nSH2, cSH2 and iSH2 (see model in **Figure 4C,D**). By disrupting the association between p110 and of the SH2 domains, the SH3 domain would activate PI3K.

## DISCUSSION

Although it was known for 30 years that certain SH3 domains can activate the PI3K pathway, the molecular basis for this activation has remained elusive. Moreover, it has been unclear how an intrinsically promiscuous SH3 association can control the activation of the PI3K pathway, of which erroneous activation can have pathological consequences for an organism. Our results now propose that the answers to both questions are linked.

Our analysis revealed two hotspot positions that modulate the PI3K activation capability of SH3 domains without affecting their interaction strength towards the canonical PR1 motif. One is the PR-proximal site, corresponding to the “R” position of the SH3 RT loop (residue 96 in Fyn), and the other is the PR-distal position located on the opposite side of the SH3 domain (residue 112 in Fyn). Our data suggest that these interactions allow PR1-bound SH3 domains to additionally associate with the C-terminal SH2 domains of p85, alleviating their inhibitory interactions on p110.

Interestingly, the strongest PI3K activation was achieved by Fyn with the PR-proximal W96, rather than the wild-type Fyn R96 which had the highest affinity for p85ΔSH3. Despite the “R” position being highly variable in SH3 domains, it is normally not a tryptophan in vertebrates. However, the SH3 domain of v-Src, the transforming gene product of the avian Rous sarcoma virus, contains a tryptophan in this position, although it derives from chicken c-Src that encodes an arginine there. Cell transformation by v-Src enhances activation of the PI3K pathway (Penuel & Martin, 1999), raising the possibility that the tryptophan-containing v-Src SH3 domain contributes to this phenotype. Interestingly, point mutations of the *FYN* gene at R96 and L112 could be detected in multiple human cancer patient samples (Sweeney et al., 2017). In particular, R96W was found in cancer of breast, colon or prostate, further raising the possibility of its role in cell transformation.

Our findings allowed us to predict the PI3K activation potential of the SH3 domains from Hck, Grb2, p85 and Ubs3a based on the stereochemistry of their PR-distal sites. For Hck, a Src kinase member and close homologue of Fyn, this activation had previously been reported (Pleiman et al., 1994). The cSH3 domain of the adaptor protein Grb2 is known to bind to p85 (Wang et al., 1995). However, it was not previously known that this binding directly activates PI3K. The capability of Grb2 to activate PI3K could allow Grb2-interacting receptors to activate PI3K, even if the receptors themselves lack the capacity to bind and activate PI3-kinase. Such a mechanism could have wide-ranging biological implications, including processes related to pathogen-infected macrophages or tumorigenesis (Haidar et al., 2015; Sastry et al., 1997; Weinger et al., 2008). The SH3 domain of p85 is known to bind to PR1 (Backer, 2010; Kapeller et al., 1994). However, this association only occurs *in trans* in the context of a p85 dimer, because the linker between the SH3 domain and PR1 is too short to allow an association *in cis* (Cheung et al., 2015). This SH3:PR1 interaction *in trans* is pivotal for p85 dimerisation and its stabilisation and activation of the PTEN phosphatase, resulting in termination of the PI3K signalling pathway (Cheung et al., 2015). However, it is only monomeric p85 that associates with p110 to form the PI3K heterodimer (Cheung et al., 2015; Layton et al., 1998). Hence, the incapability of an p85 SH3 domain to activate the PI3K pathway may help avoiding erroneous self-activation from p85:p110 complexes accumulating in the same cellular location. Finally, Ubs3a (also known as UBASH3A or STS2) regulates the localisation and activation of transmembrane receptors, including EGFR, PDGFRB and the T-cell receptor (Ge et al., 2019; Kowanetz et al., 2004). Among the actions of Ubs3A, it has been suggested that it uses its SH3 domain to sequester dynamin (Bertelsen et al., 2007). Considering that Ubs3A colocalises within the same receptor protein complexes as PI3K (Blaize et al., 2020), the non-activating SH3 association with PI3K may be biologically relevant.

Our results corroborated that PI3K is activated by SH3 domains which have only a moderate affinity for p85. This shows that the specificity of the SH3-mediated activation does not stem from an especially strong association. Consequently, PI3K could, in theory, be mistakenly activated by a variety of SH3 domains. To explain how, nonetheless, the SH3-mediated PI3K activation maintains specificity, our data suggest that p85 selects SH3 domains through function rather than through affinity. Hence, we propose that PI3K activation is controlled by “functional selectivity”; i.e. many SH3 domains can bind to the p85 PR1 motif, however PI3K activation only results from those SH3 domains capable of additional associations with the p85 nicSH2 region. Based on our data we propose that these additional interactions release the p85-imposed inhibition of p110 by competing with the association between the p85 SH2 domains and p110.

By spatially and mechanistically uncoupling “binding” and “activation”, functional selectivity can control activation in cases where erroneous binding events cannot be sufficiently excluded. Indeed, although some examples of highly specific SH3:ligand interactions exist (Seet et al., 2007), a significant cross-reactivity has been observed for SH3 domains of higher eukaryotes (Kärkkäinen et al., 2006). The membrane-associated cellular loci where PI3K is activated amidst several types of receptor-assembled signalosomes harbour many SH3-containing proteins. Our results infer that many of these nonspecific SH3 domains may bind to the PR1 region of p85, but only few SH3 domains will trigger a functional outcome using appropriate tertiary interactions.

The separation of binding and activation might also allow the creation of additional biological functions. For example, this mechanism would allow non-activating SH3 domains to recruit and possibly sequester PI3K to a subcellular environment without stimulating its enzymatic activity. Functional selectivity would also allow negative feedback loops in which an ensemble of non-activating SH3 domains out-compete activating SH3 domains. The interplay between selection through function and selection through affinity might thus enhance the ability of a cell to control specificity, function, and localisation of proteins. The proposed concept of functional selectivity synergises with other mechanisms (such as colocalization, co-expression, post-translational modifications, or multiple simultaneous interactions) to ensure selectivity in signal transduction. More generally, the concept of functional selectivity may help explain how canonical associations between ubiquitous homologous domains can achieve fidelity in signalling.

## MATERIALS AND METHODS

### Protein production and expression

The SH3 domain of Fyn, along with its R96I and R96W mutants (referred to as FynR96, FynR96I, and FynR96W respectively), spanning residues 72–142 of the human Fyn protein, Grb2 cSH3 domain (residues 151–217 of human Grb2), HCK SH3 (residues 78–138 of human HCK) were cloned into a pGEX-2T vector using BamHI-EcoRI restriction sites and expressed as GST fusion proteins. Similarly, p85 SH3 (residues 1–79 of human p85), and Ubs3A (residues 276–340 of human Ubs3a) were cloned into pGEX-6P1. All the proteins were expressed in E. coli BL21 (DE3) cells and purification was carried out as described previously (Aldehaiman et al., 2021; Arold et al., 1998).

For the full-length p85 subunit of human PI3K, p85ΔSH3 (residue 80-724), PR1-BH-PR2 (residues 80–333), and BHnicSH2 (residues 110–428), the sequences were cloned into a pGEX-6P1 vector and expressed in *E. coli* BL21 cells as GST fusion proteins. Cells were grown to an absorbance of 0.6 at 600 nm, induced with 0.2 mM IPTG, and cultured for 6 hours at 30 °C. After pelleting the cells by centrifugation, the pellets were resuspended in a buffer containing 50mM Tris-Hcl, 350mM NaCl, 2mm EDTA, 0.1% Triton X-100, 2mM β-mercaptoethanol, a pinch of lysozyme, one tablet of protease inhibitor, and stored at –80°C. For purification, the cells were thawed, then lysed using sonication. The resulting supernatant was subsequently passed through a glutathione-Sepharose column. After thorough washing with lysis buffers, the proteins were cleaved from GST by overnight treatment with 3C protease at 4 °C. The final step involved gel filtration chromatography using superdex 200 (Cytiva) in a buffer containing 20 mM hepes, 150mM NaCl, 2mM TCEP, 2 mM EDTA. The eluted peak fractions were then concentrated to 3–5 mg/ml. For NMR experiments ^15^N-labelled SH3 domains were grown in M9 medium supplemented with ^15^NH4Cl and purified as described above. The PDGFR doubly phosphorylated pY peptide G^738G^(pY)MDMSKDESVD(pY)VPML^755^, PR1 peptide (KISPPTPKPRPPRPLPVAPGS) and PR2 peptide (PAPALPPKPPKPTT) were commercially synthesised (Genscript)

### Nuclear magnetic resonance (NMR)

#### Protein expression and purification

The Fyn SH3 R96 WT and the mutants constructs of SH3 I96 and SH3 W96 were expressed in *E.coli* Bl21 (DE3) cells using expression vector with a His-tag at the C-terminal. Bacterial cells were cultured in 1 litre of M9 medium that contains 1g of ^15^N ammonium chloride and 2 g of ^13^C D-glucose at 37°C and induced at OD = 0.6 with 0.4 mM isopropyl-β-D-thio-galactoside (IPTG) and then incubated overnight at 25°C. The cells of Fyn SH3 WT and SH3 R96I were harvested and suspended in in 50 mM Tris (pH 8.0), 500 mM NaCl, 2 mM DTT, 1 tablet protease inhibitor (Roche), 1% triton and 0.4 mg/ml lysozyme lysed using mild sonication and cleared by spinning for 35 min at 28,000xg. Proteins were purified by immobilised nickel affinity column using standard procedures, followed by a Superdex 75 16/600 (GE Healthcare) size-exclusion chromatography. Fractions containing pure proteins were pooled and concentrated.

p85ΔSH3 was expressed in E-coli cells using a pGEX-6-P-1 vector with N-terminal GST tag. Bacterial cells were cultured in 1 litre of YT medium at 37°C and induced at OD = 0.6 with 0.4 mM isopropyl-β-D-thio-galactoside (IPTG) and then incubated overnight at 18°C. The cells were harvested and suspended in 50 mM Tris (pH 8.0), 200 mM NaCl, 2 mM DTT, 1 tablet protease inhibitor (Roche), 1% triton, and 0.4 mg/ml lysozyme lysed using mild sonication and cleared by spinning for 35 min at 35,000xg. Proteins were purified by immobilised GST column using standard procedures, followed by Mono Q and Superdex200 16/600 (GE Healthcare) size-exclusion chromatography. Fractions containing pure proteins were pooled and concentrated.

#### NMR samples preparation and Data recording

Protein samples were dialyzed into 20 mM sodium phosphate (pH 7.4), 150 mM NaCl, 1 mM TECP, and 0.02% Sodium Azide 90%/10% H2O/D2O. All NMR experiments were performed on a Bruker 700 MHz Avance III spectrometer equipped with the TCI 700 H-C/ N-D CryoProbe. Unless stated otherwise, experiments were measured at the temperature carefully adjusted to 25°C. Assignment of backbone ^1^H, ^15^N and ^13^C resonances was done basing on standard set of NMR experiments: 2D ^1^H,^15^N HSQC, ^1^H,^13^C HSQC spectra (Bodenhausen and Ruben, 1980) (tuned separately to aliphatic and aromatic carbons), HNCO (Muhandiram and Kay, 1994), HNCA (Ikura

et al., 1990), HNCACB (Wittekind and Mueller, 1993), CACB(CO)NH (Grzesiekt and Bax, 1992), HN(CA)CO (Clubb et al., 1992), performed on a sample containing 500uM ^13^C,^15^N-labelled Fyn I96. Assignment of Fyn WT and Fyn W96 proteins was obtained by comparison of ^1^H,^15^N HSQC spectra with assigned ^1^H,^15^N HSQC obtained for Fyn I96. Backbone connectivities were confirmed in case of both proteins by analysis of CACB(CO)NH and HNCACB. To obtain reliable assignment in 5°C we additionally measured ^1^H,^15^N HSQC and ^1^H,^13^C HSQC at 5°C, 10°C, 15°C, 20°C for Fyn I96 (with all the data collected for Fyn R96 and W96, the measurement at 5°C was sufficient to transfer assignment to lower temperature).

p85 PR1 titrations were performed by adding varying amounts of unlabeled peptide to 100uM ^13^C,^15^N-labelled Fyn R96, Fyn I96 and Fyn W96 samples respectively in ratio of (0.5, 1:1, 1:2, 1:4, 1:8, 1:16). For each titration step ^15^N,^1^H HSQC and ^13^C,^1^H HSQCs (tuned separately to aliphatic and aromatic carbons) were measured.

p85ΔSH3 titrations were performed at 5°C. Due to the low stability and solubility of p85ΔSH3 we set up the titrations in a way that we titrated increasing amounts of ^15^N,^13^C-labelled SH3 domains onto unlabeled p85ΔSH3 kept at a constant concentration. Ratios of p85ΔSH3:SH3 domains were 1:0.1, 1:0.25, 1:0.5, 1:1, 1:2, 1:3, 1:4, 1:6, 1:8, 1:10, 1:15, and 1:25 in case of each protein. We measured ^15^N,^1^H HSQC and ^13^C,^1^H HSQCs (tuned separately to aliphatic and aromatic carbons) at each titration step.

All 2D NMR experiments were processed using NMRpipe (Delaglio et al., 1995), all 3D NMR experiments were measured with sparse sampling of indirectly detected dimensions to increase the resolution and were processed using qMDD (Kazimierczuk and Orekhov, 2011; Orekhov and Jaravine, 2011) and analysed using SPARKY software (Goddard, T.D. and Kneller, D.G., SPARKY3: University of California, San Francisco.)

### ITC analysis

Isothermal titration calorimetry (ITC) experiments were performed on MicroCal PEAQ-ITC (Malvern Panalytical) and MicroCal ITC200 (GE Healthcare), using the 19-injection standard method at 25°C. All proteins were dialysed and degassed in 20 mM sodium phosphate (pH, 7.5), 150 mM NaCl, 2 mM EDTA, and 1-2 mM TCEP. For all peptide-to-protein titrations, the lyophilised peptides were dissolved in the dialysate and placed in the syringe. For titrations involving p85 domains, the SH3 domains were placed in the syringe. The protein concentrations in the cell ranged from 20 to 75 μM and were ≥10-fold higher in the syringe. The measurements and data were analysed using the software provided by Origin.

### Differential scanning fluorimetry (DSF)

All DSF experiments were performed on C1000 Touch thermal cycler CFX96 Real-Time system (Bio-Rad) with a final concentration of 50 μM SH3 with 10x Sypro orange dye (5000x Invitrogen) in 50 mM Tris (pH=8.0), 200 mM NaCl, 2 mM EDTA.

### Small angle X-ray scattering (SAXS)

SAXS measurements were performed on the purified PR1_BH_PR2 protein at the SWING beamline (SOLEIL, Saint-Aubin, France) utilising a wavelength (λ) of 1.03 Å. The sample was positioned 1.8 m away from the detector, covering a momentum transfer range of 0.01 Å−1 to 0.5 Å−1. Data acquisition was carried out via size exclusion chromatography coupled with SAXS at a temperature of 10°C, with the protein concentration maintained at 20 mg/ml. Buffer contributions were accounted for using buffer data derived from eluted buffer (20 mM HEPES, 150 mM NaCl, pH 7.5) preceding protein elution, subtracted utilising SWING’s FOXTROT software (Thureau et al., 2021). Subsequent analysis was performed employing PRIMUS and EOM modules within the ATSAS software suite (Manalastas-Cantos et al., 2021).

### Cross-linking and mass spectrometry

To initiate the crosslinking reaction, 0.1 mM bissulfosuccinimidyl suberate (BS3) was added to a mixture containing 5 µM p85ΔSH3 and 10 µM SH3 domains. The crosslinking reaction was allowed to proceed for 20 minutes at 25 °C, following which it was quenched by adding 100 mM Tris-HCl at pH 7.5. Subsequently, the cross-linked proteins were separated using a 4–12% SDS-PAGE gel. The targeted protein bands were manually excised from the SDS-PAGE gel, cut into 1 mm pieces, and subjected to washing with a solution of 100 mM ammonium bicarbonate and acetonitrile in a 1:1 ratio, followed by an additional wash in a 1:1 mixture of 25 mM ammonium bicarbonate and acetonitrile to ensure complete destaining. Subsequently, the gel pieces were treated with acetonitrile to remove residual moisture and then reduced using 10 mM DTT at 56°C for 60 minutes. The proteins within the gel pieces were then alkylated with 55 mM iodoacetamide at room temperature for 30 minutes. Afterward, the gel pieces were washed twice with a 1:1 mixture of 25 mM ammonium bicarbonate and acetonitrile to remove any excess DTT and iodoacetamide. 2 µL of a trypsin solution (1 µg/µL) in 25 mM ammonium bicarbonate was added to the gel pieces, and they were allowed to swell on ice for 60 minutes. To extract peptides, 100 µL of a solution containing 50% acetonitrile and 0.1% formic acid was added three times, and the supernatants were combined. The combined extracts were then concentrated using a speed vacuum system and resuspended in 20 µL of 0.1% formic acid. Finally, the samples were subjected to LC-MS analysis.

### LC/MS and Data processing

LC-MS/MS analysis was carried out on the Orbitrap Fusion Lumos mass spectrometer (ThermoFisher Scientific) coupled to a Dionex UltiMate 3000 RSLC nano HPLC system (Thermo Scientific). The peptide samples were loaded onto an Acclaim PepMap^TM^ 100 C18 trap column (100 µm x 2 cm, 100 A; Thermo Scientific) and resolved on a PepMap RSLC C18 analytical column (2µm, 100A, 75 µm x 50 cm; Thermo Scientific) at a flow rate of 300 nL/min over a 75 minutes gradient. Water with 0.1% FA (solvent A) and 95% ACN with 0.1% FA (solvent B) were used as mobile phases. The peptides were eluted from the column over a 55-minute multistep gradient using 2.1% to 31.6% of solvent B followed by a 5-minute gradient to 90% B. The column was then maintained at 90% B for another 3 minutes to ensure elution of all the peptides before final equilibration with 2.1% B for 10 minutes.

The spray was initiated by applying 2 kV to the EASY-Spray emitter. The data were acquired under the control of Xcalibur software in a data-dependent mode using top speed and 3 s duration per cycle. The Orbitrap was operated in positive ionisation mode, using the lock mass option (reference ion at m/z 445.120025) to ensure more accurate mass measurements in FTMS mode. The survey scan was recorded covering the *m*/*z* ranges from 400 and 1600 Th in profile mode with a resolution set to 120,000. The automatic gain control (AGC) target was set to standard and the maximum injection time was set to 50 ms. The most intense ions with charge state 2+ to 6+ were selected for fragmentation using HCD with a fixed collision energy mode. The HCD collision energy was set to 30%, and the precursor ions were isolated using a 1.6 Th window. The AGC target was set to standard with a maximum injection time mode set to dynamic. The spectra were acquired using a 15,000 resolution.

Peak lists in the format of mgf files were submitted for searching through the Protein Prospector ’Batch-Tag Web’ tool (Trnka et al., 2014). Each spectrum’s peaks were queried with a precursor charge range of 2–5, a tolerance of 20 ppm for precursor ions, and 1 Da for fragment ions. The instrument setting defined was ESI-Q-hi-res. Cleavage selectivity was adjusted to trypsin, allowing for up to two missed cleavages per peptide. Carbamidomethyl [C] and oxidation [M] were designated as constant modifications. The Protein Prospector ’Search Compare’ program was used to generate a report on Crosslinked Peptides.

### Measurement of PI3K lipid kinase activity

Endometrial cancer cells (EFE184) were lysed with immunoprecipitation lysis buffer (50 mM Tris, 150 mM NaCl, 0.5% NP-40, 5 mM ethylenediaminetetraacetic acid, protease and phosphatase inhibitors). The protein lysates were pre-cleared with protein A/G agarose beads (Santa Cruz Biotechnology, Dallas, TX) for 1 hr at 4°C prior to immunoprecipitation of p110α protein. Lysate (1 mg per reaction) was incubated with anti-p110α antibody (4255; Cell Signaling Technology, Danvers, MA) or normal rabbit IgG (Santa Cruz Biotechnology) at 1:50 dilution overnight at 4°C, followed by incubation with A/G agarose beads for 4 hr. After washing the beads, the immunoprecipitated protein was eluted by incubating with 20 μl low-pH non-denaturing glycine buffer (100 mM glycine/HCl, pH 2.5) for 10 min at room temperature. To neutralise the low pH, 2 μl 1M Tris/HCl pH 8.5 was added. The same volume of supernatant was allocated to each reaction for equal amount of immunoprecipitated p110 protein, which was then incubated with 100 μM SH3 domain peptide in the presence or absence of 2 μM PDGFR pY peptide [G^738^G(pY)MDMSKDESVD(pY)VPML^755^] at room temperature for 2 hr. Subsequently, the mixture was subjected to PI3-kinase activity assay using PI3-Kinase Activity ELISA kit (K-1000s, Echelon Biosciences, Salt Lake City, UT) according to manufacturer’s instruction. Briefly, the mixture was incubated with 50 μl PI(4,5)P2 substrate and 25 μl KBZ reaction buffer at 37 °C for 3 hr before incubation with PI(3,4,5)P3 detector for another 1 hr at room temperature. The PIP3-containing samples were then transferred to the detection Plate and incubated for 1 hr followed by 30 min incubation with the secondary detector. A signal was developed by TMB solution, and the reaction was stopped by H2SO4 stop solution. Absorbance reading was measured at 450 nm and PIP3 level was calculated by sigmoidal dose-response nonlinear regression standard curve.

### AlphaFold Structure Prediction

Modelling was done on the KAUST IBEX cluster, using an in-house wrapper (’Guzmán-Vega and ’Arold, 2024) for AlphaFold_multimer (Evans et al., 2022).

## ACKNOWLEDGEMENTS

The research reported in this publication was supported by King Abdullah University of Science and Technology (KAUST) through the baseline fund (BAS/1/1056-01) to STA and MJ. JEL acknowledges support from the Cancer Research UK Grant C57233/A22356. LWC was supported by the Hong Kong Research Grants Council (17122021, 17104022), and JWB was supported by NCI 1P01CA257885. We acknowledge SOLEIL for provision of synchrotron radiation facilities and would like to thank P. Legrand, S. Sirigu, M. Savko and B. Shepard for assistance in using the beamlines PROXIMA 1 and PROXIMA 2A, and Aurelien Thureau and Javier Perez for assistance using the beamline SWING. For computer time, this research used the resources of the KAUST Supercomputing Laboratory, and experimental research was supported by the Bioscience Core Lab, ACL Proteomics Lab and the Imaging and Characterization Core Lab at King Abdullah University of Science & Technology (KAUST) in Thuwal, Saudi Arabia.

## AUTHOR CONTRIBUTIONS

Conception of the project: STA. Recombinant protein production: SSA, AA, AS, SA, AL, STA; Experimental data acquisition and analysis for ITC: SSA, AA, AS, SA, STA, JEL; NMR: SSA, AL, XM, MN; XLMS: AA; Activity assays: HW, JB, VCYM, LWTC. Contributed reagents, instrument time and lab facilities: JB, MJ, STA, XM, LWTC. Supervision: JB, STA, JEL, MN, AA, XM. Writing of the initial manuscript: STA. All authors read and commented on the manuscript.

## SUPPLEMENTARY INFORMATION

**Supplementary Figure 1. Typical ITC titrations for data in Table 1**. All plots show raw heats (adjusted to baseline) on top, and integrated heats below. All titrations were carried out in triplicates.

**Supplementary Figure 2. HSQC plots for titrations underlying Figure 3A**. Overlay of ^1^H-^15^N HSQC spectra acquired during p85 PR1 titrations. Varying amounts of unlabelled peptide were added to 100 µM ^13^C,^15^N-labelled Fyn R96, Fyn I96, and Fyn W96 samples.

**Supplementary Figure 3. PR1-BH-PR2 SAXS data.** Ensemble Optimization Method (EOM) analysis of SEC-SAXS data collected on the p85 PR1-BH-PR2 fragment. The BH domain (residues 121-299) was kept fixed, and the flanking PR1 and PR2 regions were allowed to be flexible. The radius of gyration (Rg; **A**) and maximum distance (Dmax; **B**) are shown in Å for the structural pool (black) and the EOM selected representative structures (red). **C**) Fit of selected ensemble models (red line) to SAXS data (black dots) with chi^2^ of 7.42. **D**) EOM-selected structures are shown superimposed on their BH domain (green). Collectively, the distributions of the selected molecules from the Rg and Dmax pools, and the conformations of the PR1 and PR2 regions in the structural models support that both PR regions are highly flexible and do not stably associate with the BH domain.

**Supplementary Figure 4. Differential scanning fluorimetry to determine the thermal stability of 50 µM Fyn SH3 domains** (wild-type and mutants). **A**) Melting curve. **B**) First derivative of (A). Traces for the SH3 are colour-coded as in the figure legends. **C**) Table showing the resulting melting temperature, *Tm*, in °C.

**Supplementary Figure 5. Additional figures for XLMS analysis shown in Figure 4B**. A Coomassie-stained SDS-PAGE gel illustrates the crosslinking of the ΔSH3 with SH3 domains. In Lane 1, FynSH3 is crosslinked with ΔSH3; in Lane 2, Grb2 cSH3 is crosslinked with ΔSH3; and in Lane 3, p85SH3 is crosslinked with ΔSH3. The full-length p85 was included as a reference for the size of crosslinked proteins. Bands highlighted within red boxes were excised from the gel, then subjected to trypsin digestion, and subsequently analysed by LC/MS.

Supplementary Figure 6. **Additional figures for XLMS analysis shown in Figure 4B**. List of Inter-protein crosslinks detected in protein prospector analysis from the LC/MS data. Crosslinks between ΔSH3 and A) p85SH3; B) Fyn SH3; C) Grb2 cSH3.

**Supplementary Figure 7. Additional figures for modelling analysis. A**) Superimposition of the AlphaFold model of the p110:p85 complex (colours as in Figure 4B), with the experimental structure PDB 8DCP (p110: light grey; p85: pale cyan). The panels show the membrane view (*left*) and back view (*right*). The SH domains of p85 are indicated. **B**) Predicted Aligned Error (PAE) plot for the AlphaFold prediction of the p110:p85 complex. p110 was given as sequence A and p85 as sequence B. The regions corresponding to the nSH2, iSH2 and cSH2 domains are indicated. **C**) Detail of the AlphaFold model of the p110:p85 complex. The p110 catalytic domain is shown as a grey surface. p85 is shown in ribbon presentation with iSH2 in magenta, cSH2 in yellow, and the linker in grey. The Y^607^SLV motif of the linker contacting the p110 catalytic domain is highlighted as a stick model with carbon atoms in orange.

