## Supplementary information for "Functional selection in SH3-mediated activation of the PI3 kinase"

1 Biological and Environmental Science and Engineering Division, Computational Biology Research Center, King Abdullah University of Science and Technology (KAUST), Thuwal 23955-6900, Kingdom of Saudi Arabia.
2 School of Biomedical Sciences, Li Ka Shing Faculty of Medicine, The University of Hong Kong, Hong Kong, China
3 Department of Molecular Pharmacology, Albert Einstein College of Medicine, Bronx, New York, USA
4 CRCM, CNRS, INSERM, Institut Paoli-Calmettes, Aix-Marseille University, 13009 Marseille, France
5 School of Molecular and Cellular Biology, and Astbury Centre for Structural Molecular Biology, University of Leeds, Leeds, LS2 9JT,
6 Faculty of Chemistry, Biological and Chemical Research Centre, University of Warsaw, Żwirki i Wigury 101, 02-089 Warsaw, Poland
* contributed equally
% Present Address: Infectious Disease Research Department, King Abdullah International Medical Research Center, Riyadh, Saudi Arabia
& Present Address: King Saud bin Abdulaziz University for Health Science, Riyadh, Saudi Arabia
! Present Address: Sartorius Canada Inc., 1173 North Service Rd W D4, L6M2V9 Oakville, ON, Canada

List of contents

Supplementary Figure 1
Supplementary Figure 2
Supplementary Figure 3
Supplementary Figure 4
Supplementary Figure 5
Supplementary Figure 6
Supplementary Figure 7

### Supplementary Figure 1

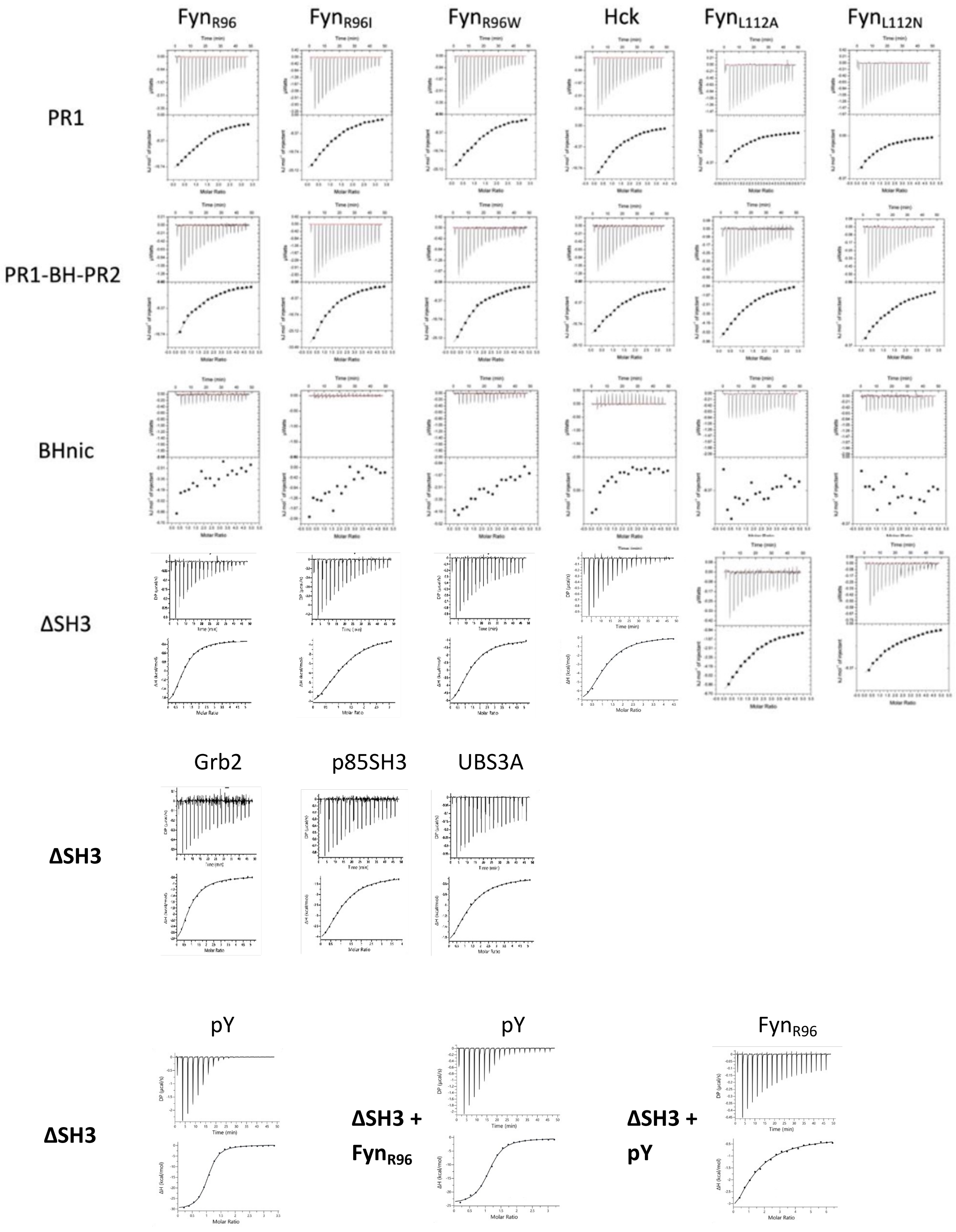


**Supplementary Figure 1. Typical ITC titrations for data are in Table 1.** All plots show raw heats (adjusted to baseline) on top, and integrated heats below. All titrations were carried out in triplicates.

Supplementary Figure 2

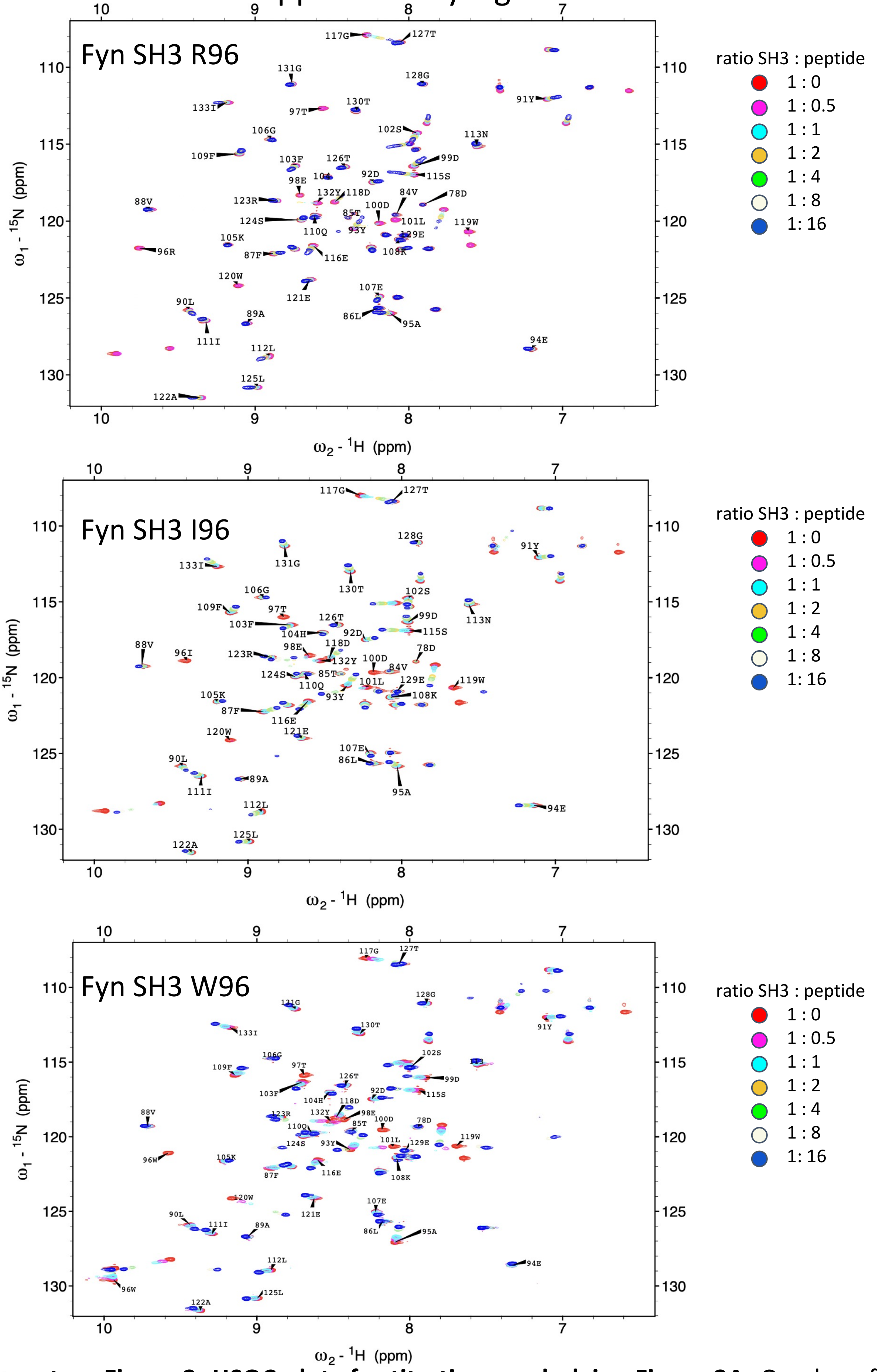


**Supplementary Figure 2. HSQC plots for titrations underlying Figure 3A.** Overlay of $^{1}H$-$^{15}N$ HSQC spectra acquired during p85 PR1 titrations. Varying amounts of unlabelled peptide were added to 100 μM $^{13}C$,$^{15}N$-labelled Fyn R96, Fyn I96, and Fyn W96 samples.

### Supplementary Figure 3

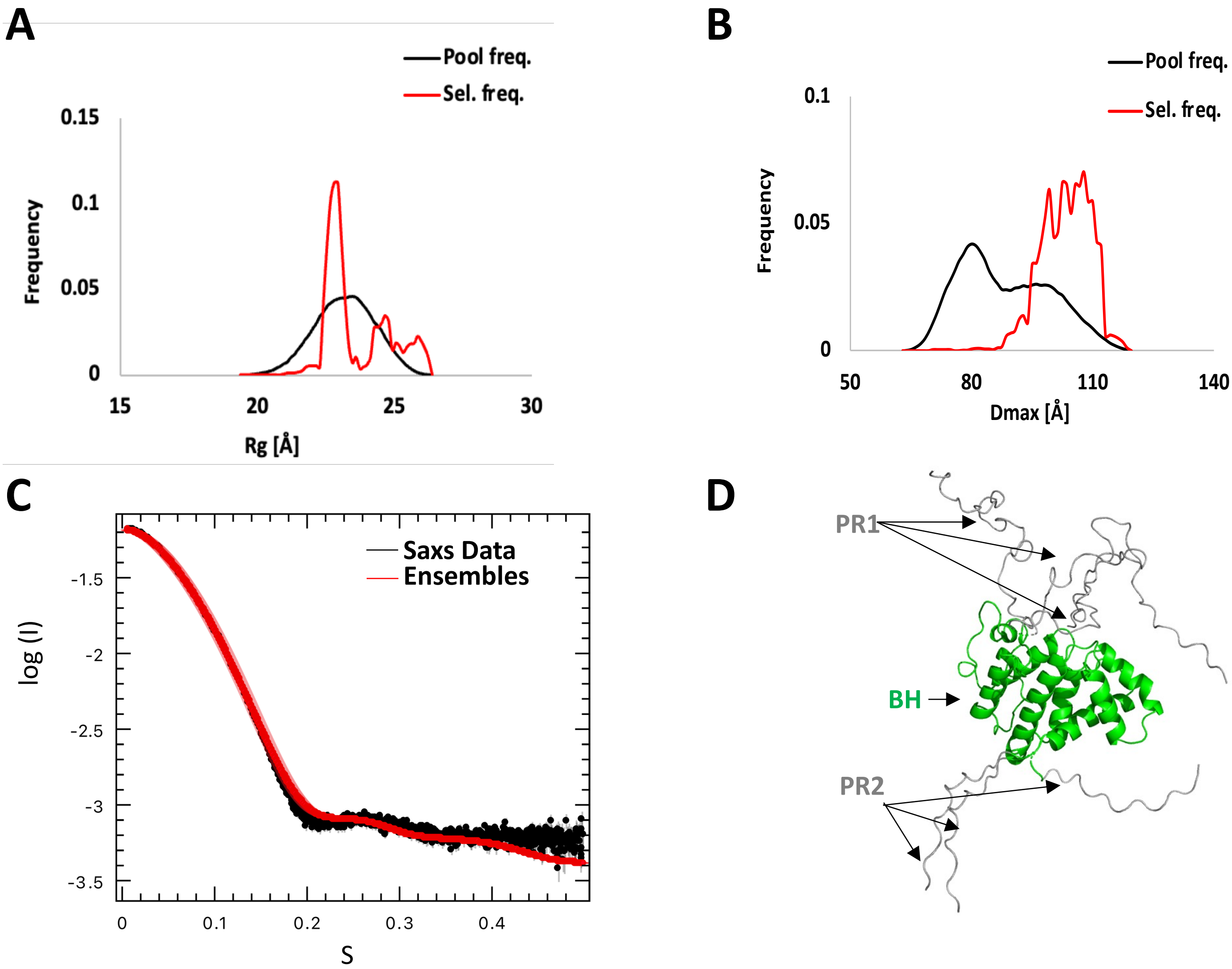


**Supplementary Figure 3. PR1-BH-PR2 SAXS data.** Ensemble Optimization Method (EOM) analysis of SEC-SAXS data collected on the p85 PR1-BH-PR2 fragment. The BH domain (residues 121-299) was kept fixed, and the flanking PR1 and PR2 regions were allowed to be flexible. The radius of gyration (Rg; **A**) and maximum distance (Dmax; **B**) are shown in Å for the structural pool (black) and the EOM selected representative structures (red). **C**) Fit of selected ensemble models (red line) to SAXS data (black dots) with $chi^2$ of 7.42. **D**) EOM-selected structures are shown superimposed on their BH domain (green). Collectively, the distributions of the selected molecules from the Rg and Dmax pools, and the conformations of the PR1 and PR2 regions in the structural models support that both PR regions are highly flexible and do not stably associate with the BH domain.

### Supplementary Figure 4

**A**

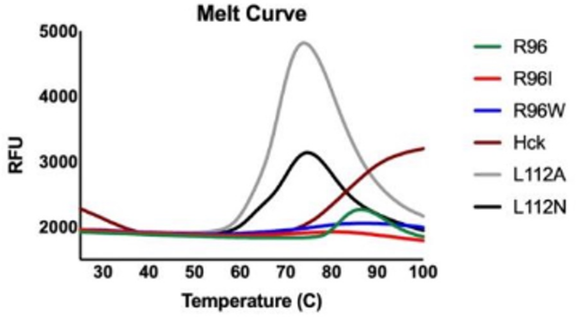


**B**

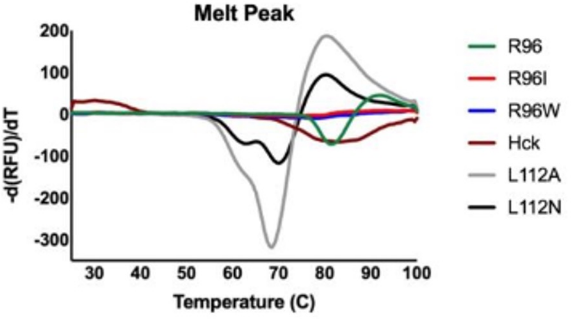


**C**

| SH3 domain Protein | Melting Temp. (C) *Tm* (˚C) |
|---|---|
| R96 | 81.8±0.3 |
| R96I | 72.5±0.5 |
| R96W | 77.5±0.5 |
| Hck | 88.5±0.5 |
| L112A | 68.0±0.5 |
| L112N | 69.7±0.6 |

**Supplementary Figure 4. Differential scanning fluorimetry to determine the thermal stability of 50 μM Fyn SH3 domains** (wild-type and mutants). **A**) Melting curve. **B**) First derivative of (A). Traces for the SH3 are colour-coded as in the figure legends. **C**) Table showing the resulting melting temperature, *Tm*, in ˚C.

### Supplementary Figure 5

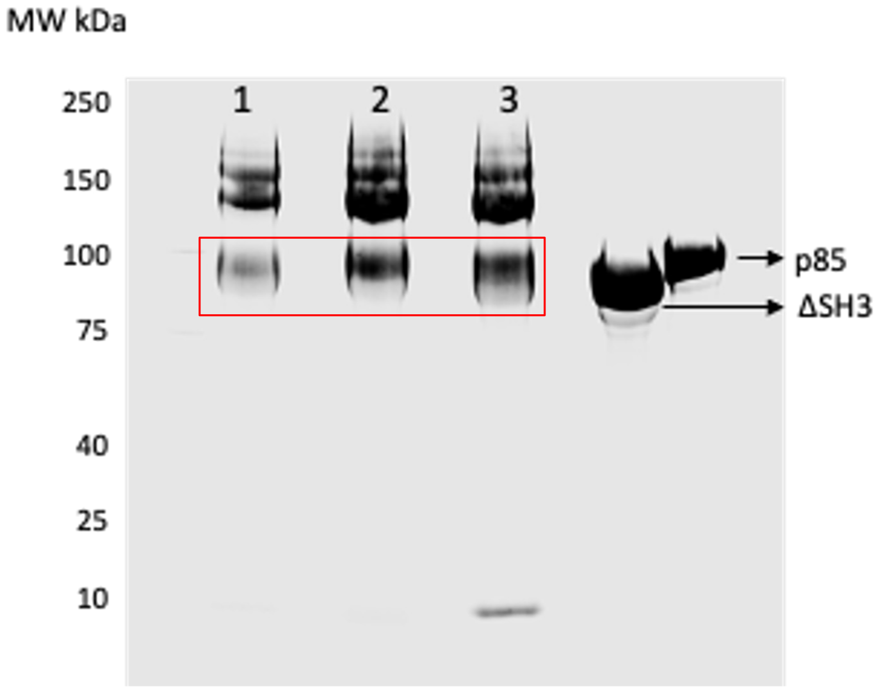


**Supplementary Figure 5. Additional figures for XLMS analysis shown in Figure 4B**. A Coomassie-stained SDS-PAGE gel illustrates the crosslinking of the ΔSH3 with SH3 domains. In Lane 1, FynSH3 is crosslinked with ΔSH3; in Lane 2, Grb2 cSH3 is crosslinked with ΔSH3; and in Lane 3, p85SH3 is crosslinked with ΔSH3. The full-length p85 was included as a reference for the size of crosslinked proteins. Bands highlighted within red boxes were excised from the gel, then subjected to trypsin digestion, and subsequently analysed by LC/MS.

Supplementary Figure 6

**A**

Inter-protein crosslinks between p85 SH3 and ΔSH3

| XLink AA 1 | peptide 1 | Protein1 | XLink AA 2 | DB Peptide 2 | Protein 2 |
|---|---|---|---|---|---|
| 0 | GPLGSEGMSAEGYQYR | pSH3 | 491 | MNSIKPDLIQLR | delSH3p85 |
| 0 | GPLGSEGMSAEGYQYR | pSH3 | 347 | LDVKLLYPVSK | delSH3p85 |
| 0 | GPLGSEGMSAEGYQYR | pSH3 | 557 | NKAENLLR | delSH3p85 |
| 0 | GPLGSEGMSAEGYQYR | pSH3 | 499 | KTR | delSH3p85 |
| 0 | GPLGSEGMSAEGYQYR | pSH3 | 0 | GPLGSISPPTPKPRPPR PLPVAPGSSK | delSH3p85 |
| 0 | GPLGSEGMSAEGYQYR | pSH3 | 0 | GPLGSISPPTPKPRPPR PLPVAPGSSK | delSH3p85 |
| 0 | GPLGSEGMSAEGYQYR | pSH3 | 303 | GGNNKLIKIFHR | delSH3p85 |
| 0 | GPLGSEGMSAEGYQYR | pSH3 | 565 | GKR | delSH3p85 |
| 0 | GPLGSEGMSAEGYQYR | pSH3 | 454 | IMHNYDKLK | delSH3p85 |
| 0 | GPLGSEGMSAEGYQYR | pSH3 | 474 | LEEDLKK | delSH3p85 |
| 0 | GPLGSEGMSAEGYQYR | pSH3 | 499 | KTR | delSH3p85 |
| 517 | KLNEWLGNENTEDQYSLVE DDEDLPHHDEK | delSH3p85 | 23 | KER | pSH3 |
| 0 | GPLGSEGMSAEGYQYR | pSH3 | 0 | GPLGSISPPTPKPRPPR PLPVAPGSSK | delSH3p85 |
| 0 | GPLGSEGMSAEGYQYR | pSH3 | 437 | FKR | delSH3p85 |
| 511 | DQYLMWLTQKGVR | delSH3p85 | 0 | GPLGSEGMSAEGYQY R | pSH3 |
| 0 | GPLGSEGMSAEGYQYR | pSH3 | 343 | NESLAQYNPKLDVK | delSH3p85 |
| 0 | GPLGSEGMSAEGYQYR | pSH3 | 454 | IMHNYDKLK | delSH3p85 |
| 0 | GPLGSEGMSAEGYQYR | pSH3 | 437 | FKR | delSH3p85 |
| 491 | MNSIKPDLIQLR | delSH3p85 | 22 | ALYDYKK | pSH3 |
| 0 | GPLGSEGMSAEGYQYR | pSH3 | 456 | IMHNYDKLKSR | delSH3p85 |
| 0 | GPLGSEGMSAEGYQYR | pSH3 | 485 | EIDKR | delSH3p85 |
| 0 | GPLGSEGMSAEGYQYR | pSH3 | 149 | KLIR | delSH3p85 |
| 22 | ALYDYKK | pSH3 | 430 | YSKEYIEK | delSH3p85 |
| 0 | GPLGSEGMSAEGYQYR | pSH3 | 475 | KQAAEYR | delSH3p85 |
| 0 | GPLGSEGMSAEGYQYR | pSH3 | 485 | EIDKR | delSH3p85 |
| 0 | GPLGSEGMSAEGYQYR | pSH3 | 180 | LSQTSSKNLLNAR | delSH3p85 |
| 443 | EGNEKEIQR | delSH3p85 | 22 | ALYDYKK | pSH3 |
| 287 | DASTKMHGDYTLTLR | delSH3p85 | 22 | ALYDYKK | pSH3 |
| 0 | GPLGSEGMSAEGYQYR | pSH3 | 485 | EIDKR | delSH3p85 |
| 0 | GPLGSEGMSAEGYQYR | pSH3 | 287 | DASTKMHGDYTLTLR | delSH3p85 |
| 0 | GPLGSEGMSAEGYQYR | pSH3 | 474 | RLEEDLKK | delSH3p85 |
| 557 | NKAENLLR | delSH3p85 | 22 | ALYDYKK | pSH3 |
| 491 | MNSIKPDLIQLR | delSH3p85 | 23 | KER | pSH3 |
| 443 | EGNEKEIQR | delSH3p85 | 23 | KER | pSH3 |

**B** Inter-protein crosslinks between Fyn SH3 and ΔSH3

| XLink AA 1 | Peptide 1 | Protein1 | XLink AA 2 | Peptide 2 | Protein2 |
|---|---|---|---|---|---|
| 28 | TEDDLSFHKGEKFQILNSSEG DWWEAR | Fyn | 343 | NESLAQYNPKLDVK | delSH3p85 |
| 25 | TEDDLSFHKGEKFQILNSSEG DWWEAR | Fyn | 475 | LEEDLKKQAAEYR | delSH3p85 |
| 28 | TEDDLSFHKGEKFQILNSSEG DWWEAR | Fyn | 435 | YSKEYIEKFK | delSH3p85 |
| 25 | TEDDLSFHKGEKFQILNSSEG DWWEAR | Fyn | 557 | NKAENLLR | delSH3p85 |
| 28 | GEKFQILNSSEGDWWEAR | Fyn | 443 | EGNEKEIQR | delSH3p85 |
| 28 | GEKFQILNSSEGDWWEAR | Fyn | 557 | NKAENLLR | delSH3p85 |
| 28 | TEDDLSFHKGEKFQILNSSEG DWWEAR | Fyn | 443 | REGNEKEIQR | delSH3p85 |
| 28 | TEDDLSFHKGEKFQILNSSEG DWWEAR | Fyn | 565 | GKR | delSH3p85 |
| 28 | TEDDLSFHKGEKFQILNSSEG DWWEAR | Fyn | 499 | KTR | delSH3p85 |
| 28 | GEKFQILNSSEGDWWEAR | Fyn | 499 | KTR | delSH3p85 |
| 0 | MTGVTLFVALYDYEAR | Fyn | 435 | EYIEKFK | delSH3p85 |
| 0 | MTGVTLFVALYDYEAR | Fyn | 474 | RLEEDLKK | delSH3p85 |

**C** Inter-protein crosslinks between Grb2 cSH3 and ΔSH3

| XLink AA 1 | Peptide 1 | Protein 1 | XLink AA 2 | Peptide 2 | Protein 2 |
|---|---|---|---|---|---|
| 413 | TAIEAFNETIKIFEEQCQTQER | p85delSH3 | 157 | KDAERQLLSFGNPR | Grb2 |
| 440 | EGNEKEIQR | p85delSH3 | 7 | MGCVQCKDKEATK | Grb2 |
| 488 | MNSIKPDLIQLR | p85delSH3 | 9 | DKEATK | Grb2 |
| 359 | LLYPVSKYQQDQVVKEDNIEAVGK | p85delSH3 | 9 | MGCVQCKDKEATK | Grb2 |
| 562 | GKRDGTFLVR | p85delSH3 | 0 | MGCVQCK | Grb2 |
| 472 | KQAAEYR | p85delSH3 | 0 | MGCVQCK | Grb2 |
| 359 | YQQDQVVKEDNIEAVGK | p85delSH3 | 322 | KLKHDK | Grb2 |
| 209 | FSAASSDNTENLIKVIEILISTEWNER | p85delSH3 | 204 | GDHVKHYKIR | Grb2 |
| 488 | MNSIKPDLIQLR | p85delSH3 | 0 | MSAEGYQYR | Grb2 |
| 488 | MNSIKPDLIQLR | p85delSH3 | 16 | KER | Grb2 |
| 427 | YSKEYIEK | p85delSH3 | 16 | KER | Grb2 |
| 488 | MNSIKPDLIQLR | p85delSH3 | 16 | KER | Grb2 |
| 413 | TAIEAFNETIKIFEEQCQTQER | p85delSH3 | 0 | MEAIAKYDFK | Grb2 |
| 195 | GDFIHVMDNSDPNWWKGACHGQTGMFPR | Grb2 | 482 | EIDKR | p85delSH3 |
| 359 | YQQDQVVKEDNIEAVGK | p85delSH3 | 0 | MEAIAK | Grb2 |
| 369 | KLHEYNTQFQEK | p85delSH3 | 10 | YDFKATADDELSFK | Grb2 |
| 195 | GDFIHVMDNSDPNWWKGACHGQTGMFPR | Grb2 | 496 | KTR | p85delSH3 |
| 63 | KGLECSTLYR | p85delSH3 | 117 | DGAGKYFLWVVK | Grb2 |
| 284 | DASTKMHGDYTLTLR | p85delSH3 | 0 | MEAIAKYDFK | Grb2 |
| 488 | RMNSIKPDLIQLR | p85delSH3 | 44 | AELNGKDGFIPK | Grb2 |
| 471 | RLEEDLKK | p85delSH3 | 0 | MEAIAK | Grb2 |
| 359 | YQQDQVVKEDNIEAVGK | p85delSH3 | 20 | YDFKATADDELSFKR | Grb2 |
| 310 | DGKYGFSDPLTFSSVVELINHYR | p85delSH3 | 6 | MEAIAKYDFK | Grb2 |
| 303 | LIKIFHR | p85delSH3 | 20 | ATADDELSFKRGDILK | Grb2 |
| 0 | KKISPPTPKPRPPRPLPVAPGSSK | p85delSH3 | 6 | MEAIAKYDFK | Grb2 |

**Supplementary Figure 6. Additional figures for XLMS analysis are shown in Figure 4B**. List of Inter-protein crosslinks detected in protein prospector analysis from the LC/MS data. Crosslinks between ΔSH3 and A) p85SH3; B) Fyn SH3; C) Grb2 cSH3.

Supplementary Figure 7

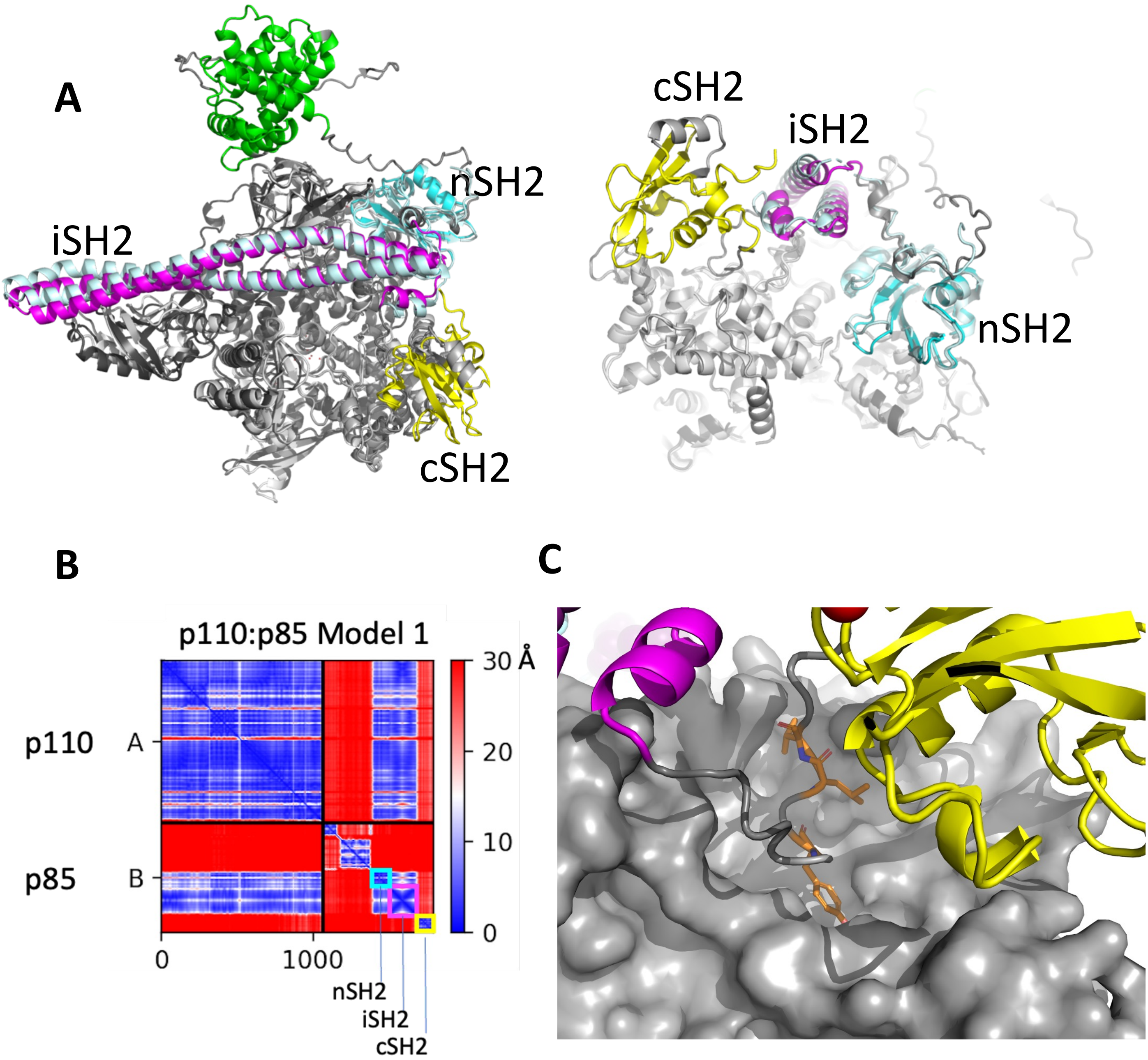


**Supplementary Figure 7. Additional figures for modelling analysis. A**) Superimposition of the AlphaFold model of the p110:p85 complex (colours as in Figure 4B), with the experimental structure PDB 8DCP (p110: light grey; p85: pale cyan). The panels show the membrane view (*left*) and back view (*right*). The SH domains of p85 are indicated. **B**) Predicted Aligned Error (PAE) plot for the AlphaFold prediction of the p110:p85 complex. p110 was given as sequence A and p85 as sequence B. The regions corresponding to the nSH2, iSH2 and cSH2 domains are indicated. **C**) Detail of the AlphaFold model of the p110:p85 complex. The p110 catalytic domain is shown as a grey surface. p85 is shown in ribbon presentation with iSH2 in magenta, cSH2 in yellow, and the linker in grey. The $Y^{607}$SLV motif of the linker contacting the p110 catalytic domain is highlighted as a stick model with carbon atoms in orange.
